# Carbon-negative biosynthesis of pyrone and pyridine dicarboxylic acids from terephthalic acid via continuous mixotrophic gas fermentation in *Cupriavidus necator* H16

**DOI:** 10.64898/2026.03.12.709797

**Authors:** Ewan Waters, Alex Conradie, Rajesh Reddy Bommareddy

## Abstract

Industrial defossilisation requires carbon-negative routes for sustainable chemical production. Bioprocesses based on renewable feedstocks are often constrained by biogenic CO₂ emissions, reducing yields and undermining environmental performance. Here, we report a mixotrophic gas fermentation strategy using the chemolithoautotroph *Cupriavidus necator* H16 that simultaneously assimilates CO₂ for cell growth and biocatalyst generation while converting the PET monomer terephthalic acid (TPA) into value-added biopolymer precursors. This process achieved complete conversion of TPA to 2-pyrone-4,6-dicarboxylic acid (PDC) with titres of 24.5 g/L and a productivity of 0.47 g/L·h. Conversion to pyridine dicarboxylic acids (2,4- and 2,5-PDCA) was less efficient (∼22% and ∼4% respectively) due to metabolic limitations such as intermediate toxicity, pH and ammonia-dependent spontaneous cyclisation. Our results establish the first carbon-negative route coupling simultaneous CO₂ assimilation with plastic monomer valorisation, providing a blueprint for sustainable biomanufacturing aligned with global climate and circular economy goals.

## Background

The transition toward carbon-neutral and carbon-negative chemical manufacturing demands bioprocesses capable of utilising sustainable, non-fossil feedstocks while simultaneously reducing greenhouse gas emissions. Gas fermentation has emerged as a compelling technology to meet this need, enabling biological conversion of waste gases such as CO₂ and CO into chemicals and fuels. Demonstrated industrial-scale deployments using anaerobic acetogens for the production of ethanol and isopropanol have validated the viability of gas fermentation as a carbon-refining platform [1, 2]. However, anaerobic systems remain constrained by metabolic limitations that restrict their accessible product spectrum, particularly for higher-value or structurally complex molecules.

Aerobic chemolithoautotrophs, most notably *Cupriavidus necator* H16, offer a more flexible alternative due to their broad metabolic capacity, robust genetic toolbox, and ability to grow solely on CO₂ and H₂ [3]. Through the Calvin–Benson–Bassham cycle, *C. necator* can fully decouple carbon assimilation from organic feedstocks, relying on renewable hydrogen for reducing power. Recent technoeconomic analyses and process innovations, including heat-integrated gas fermentation processes have demonstrated the feasibility of aerobic CO₂-to-chemical manufacturing using *C. necator* [4, 5]. Nonetheless, the high reducing-power demand associated with CO₂ fixation limits yields and productivities for pathways requiring extensive redox transformations. Mixotrophic cultivation, two-stage growth–production strategies, and alternative co-feeding regimes have been proposed to overcome these constraints, but efficient production of complex value-added chemicals from CO₂ remains a challenge[6, 7].

Parallel to the development of CO₂-based bioprocesses, valorisation of plastic waste has become an essential component of circular manufacturing. Poly(ethylene terephthalate) (PET) is one of the most widely used plastics globally, and its chemical or enzymatic depolymerisation yields terephthalic acid (TPA), a stable aromatic monomer amenable to microbial conversion[8–11]. Increasing attention has turned toward biological upcycling of TPA and other lignin- or PET-derived aromatics into polyester monomers such as 2-pyrone-4,6-dicarboxylic acid (PDC) and pyridine dicarboxylic acids (2,4-PDCA and 2,5-PDCA) [12–16]. These pseudo-aromatic dicarboxylic acids exhibit structural similarity to petroleum-derived monomers, offering potential applications in engineering polymers and functional materials. While promising, existing microbial production routes depend heavily on heterotrophic hosts and reduced organic feedstocks such as sugars or lignin-derived compounds—limiting both sustainability and cost-effectiveness. Recent works demonstrated high-level heterotrophic production of PDC and 2,4-PDCA from TPA derived from PET using engineered *P. putida* and multiple engineered *E. coli* strains respectively, achieving high titres (129 g/L) and productivities (1.67 g/L·h)[17–19]. These approaches use glucose for supporting the growth where the intrinsic problem of biogenic CO_2_ evolution still exists and implementing multi-strain systems increases process complexity.

A major opportunity exists to unite these two sustainability goals, CO₂ utilisation and plastic upcycling into a single integrated bioprocess. *C. necator* H16 is an ideal candidate for such an approach due to its capacity for high-cell-density autotrophic growth on CO₂/H₂ and its amenability to expressing heterologous aromatic ring-cleavage pathways. The conversion of TPA to pseudo-aromatic dicarboxylic acids proceeds through protocatechuic acid (PCA) and its meta-cleavage intermediates 4- and 5-carboxy-2-hydroxymuconate semialdehyde (4- and 5-CHMS) (Figure 1). These intermediates are chemically unstable and highly toxic, particularly in gram-negative hosts, often resulting in low titres when producing PDCAs [14, 15]. In contrast, PDC production proceeds through the closed CHMS form and bypasses the ammonolysis step that drives PDCA-associated toxicity, suggesting that *C. necator* may be uniquely suited for efficient PDC synthesis.

**Figure 1:**
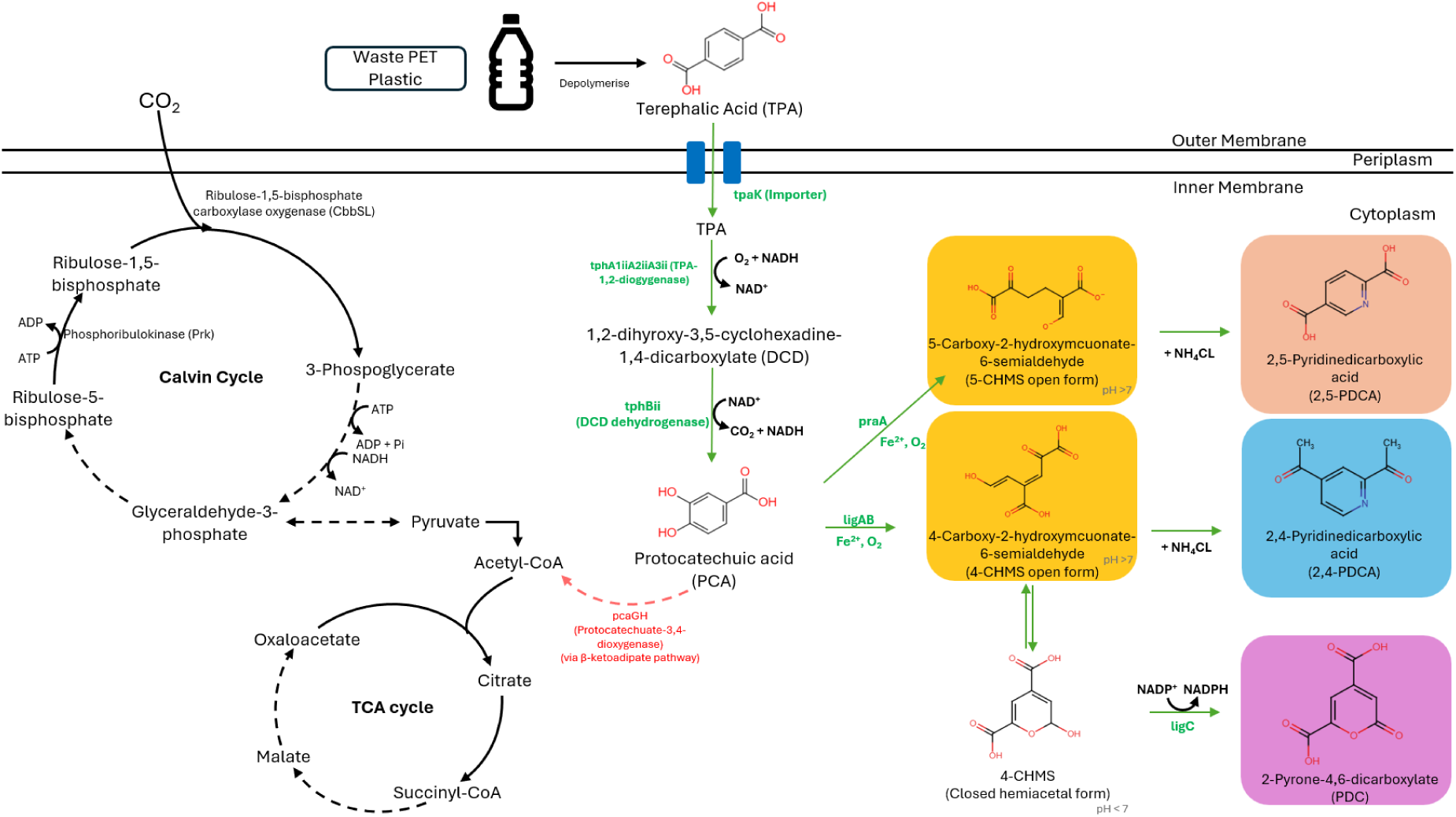
Production of PDCAs and PDC in *C. necator* H16. Autotrophic growth of *C. necator* H16 is enabled by CO_2_ assimilation into the Calvin Cycle. Simultaneously, strains can be engineered for the biocatalysis of organic carbon. The biosynthetic conversion of TPA to PDCAs and PDC is detailed with reactions requiring heterologous gene expression illustrated in green. Areas of potential carbon flux diversion into the central metabolism are illustrated in red.

Despite the rapidly growing interest in both gas fermentation and plastic upcycling, no study has yet demonstrated a unified bioprocess in which CO₂ and H₂ drive biomass formation while TPA is simultaneously converted into PDC or PDCAs in a single organism. Achieving such integration would represent a significant advancement toward carbon-negative, resource-efficient biomanufacturing, transforming two widely available waste streams such as CO₂ and PET monomers into high-value polymer precursors.

In this work, we address this gap by engineering *C. necator* H16 to co-assimilate CO₂ and bio convert TPA into three valuable dicarboxylic acids: PDC, 2,4-PDCA and 2,5-PDCA. Using metabolic engineering, pH- and buffer-controlled pathway optimisation, and both resting-cell and continuous mixotrophic gas fermentations, we establish a process that achieves near-quantitative conversion of TPA to PDC and significantly improves PDCA yields. This represents the first demonstration of a carbon-negative route that integrates CO₂ assimilation with PET-derived aromatic upcycling in a single bioprocess, providing a blueprint for future circular and sustainable manufacturing systems.

## Results

### Establishing TPA uptake and conversion to PDCAs in *C. necator* H16

*C. necator* H16 is unable to assimilate TPA but can utilise the intermediate metabolite PCA as a precursor to succinyl-CoA and acetyl-CoA via the β-ketoadipate pathway (Figure 2e). Therefore, establishing the bioconversion of TPA to dicarboxylic acid product require heterologous gene expression that would facilitate the import of TPA and the meta-cleavage of the PCA intermediate towards the desired product.

To engineer the metabolic pathway producing pyridine dicarboxylic acids from TPA (Figure 2a and b), genes belonging to the TAPDO operon (*tpaK, tphA2II, tphA3II, tphBII, tphAII from Comamonas sp E6),* and either the protocatechuate-4,5-dioxygenase, *ligAB* (*Sphingobium* sp. SYK-6) or the protocatechuate-2,3-dioxygenase, *praA* (*Paenibacillus* sp. JJ-1b) were assembled into a pBBR expression vector (Figure 2c and d) (Table S4 and S6). The codon optimised sequences are shown in the supplementary Table S5. These genes on the pBBR expression vector were all expressed under the L-arabinose inducible promoter, designated as H16 TAPDO *ligAB* and H16 TAPDO *praA*.

**Figure 2:**
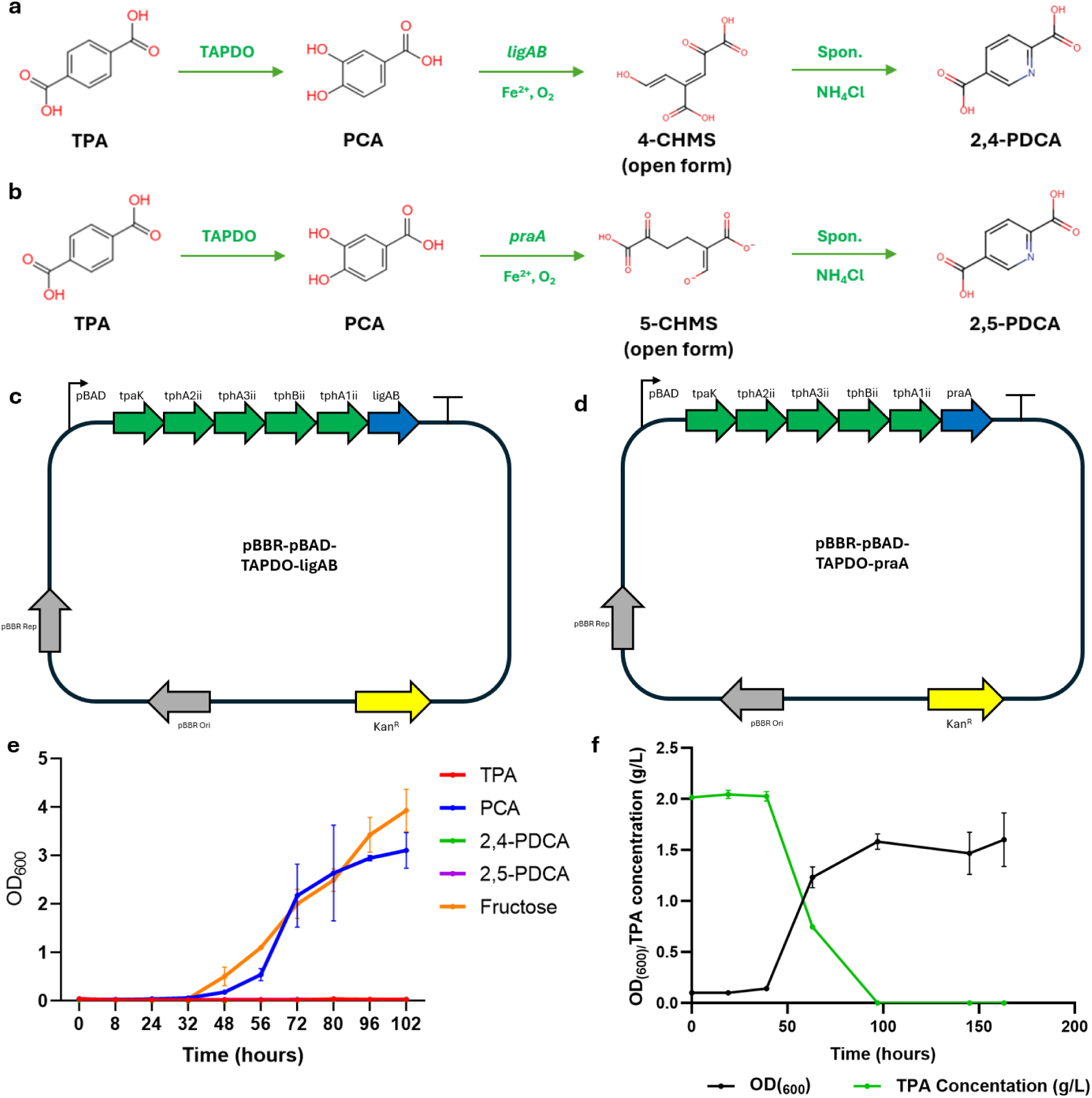
Establishing TPA biocatalysis in *C. necator* H16. Biosynthetic pathways of **a** 2,4-PDCA and **b** 2,5-PDCA production from TPA are enabled by the expression of non-native genes (green). Expression of the TPA converting TAPDO operon (green) and production genes (blue) in a pBBR plasmid facilitate production of **c** 2,4-PDCA and **d** 2,5-PDCA. **e** H16 wild-type is not capable of growth on TPA, 2,4-PDCA or 2,5-PDCA. H16 can grow on pathway intermediate protocatechuic acid (PCA) via the β-ketoadipate pathway. **f** Expression of the TAPDO operon in expression plasmids **c** and **d** facilitated the growth of H16 in minimal media supplemented with 2 g/L TPA as a sole carbon source (starting OD_600_ 0.1) The uptake of TPA (green line) is plotted alongside OD_600_ (black line) All values are a mean average of 3 replicates ± SD

As expected, expression of the TAPDO operon in H16 enabled the import of TPA that would be utilised for growth when 2 g/L TPA was used as a sole carbon source (Figure 2f). However, this bioconversion at a low optical density did not result in PDCA production despite higher expression of *ligAB* and *praA* (Figure S1). As a result, future biotransformations were carried out at OD_600_ 40 to harness a higher density of expressed heterologous proteins.

### Optimising TPA conversion to PDCAs in resting cell experiments

Once the TPA uptake has been confirmed, in the TAPDO expressing strains, subsequent work focused on optimising the conversion of TPA towards PCA and then to the PDCAs. The toxicity of the PCA-meta cleavage product 4- or 5-carboxy-2-hydroxymuconate-semialdehyde (4- and 5-CHMS) has been well documented and as previously described using resting cells for biotransformation decouples growth related inhibition from toxic chemicals[14]. H16 TAPDO *ligAB* and H16 TAPDO *praA* expressing respective operons were separately grown in LB medium and induced for protein expression at the beginning of growth for 16 hours before being washed and resuspended in resting cell buffer (PBS, 5 g/L NH_4_Cl, 2 g/L TPA, pH 7.5, OD_600_ 40). After 24 hours product conversion efficiencies were low (Table 1), with 0.31 g/L 2,4-PDCA produced and no 2,5-PDCA was detected.

**Table 1:** Bioproduction of PDCAs from TPA by H16 resting cells. The yield of PDCAs and molar conversion from 2 g/L TPA by H16 TAPDO *ligAB* and H16 TAPDO *praA* (PBS, 5 g/L NH_4_Cl, pH 7.5, OD_600_ 40, protein expression = 16 hours). Yields are means of triplicates ± SD. Quantification of metabolites was carried out by HPLC analysis

| Product | Buffer | Titre after 24 hrs | Conversion |
| --- | --- | --- | --- |
| 2,4-PDCA | PBS | 0.31 ± 0.004 g/L | 15.5% |
| 2,5-PDCA | PBS | 0.00 g/L | - |

In both reactions, colour change characteristic of 4- and 5-CHMS production was observed with a vivid yellow colour appearing minutes into the reaction (Figure 3a, 3b and 4a). CHMS detection signifies enzymatic activity of both the TAPDO operon and the ring-cleavage dioxygenases (*ligAB/praA*). Therefore, any bottleneck in yield lies within the spontaneous cyclisation of 4- or 5-CHMS to the final PDCA product. CHMS is known to accumulate in a pH-dependent equilibrium in its closed form in acidic conditions and open form in basic pH media. The open form of CHMS is yellow in colour and required for 2,4- and 2,5-PDCA production in the presence of ammonium. Consequently, efforts were undertaken to increase the accumulation of open form CHMS and improve its conversion to PDCA product.

**Figure 3:**
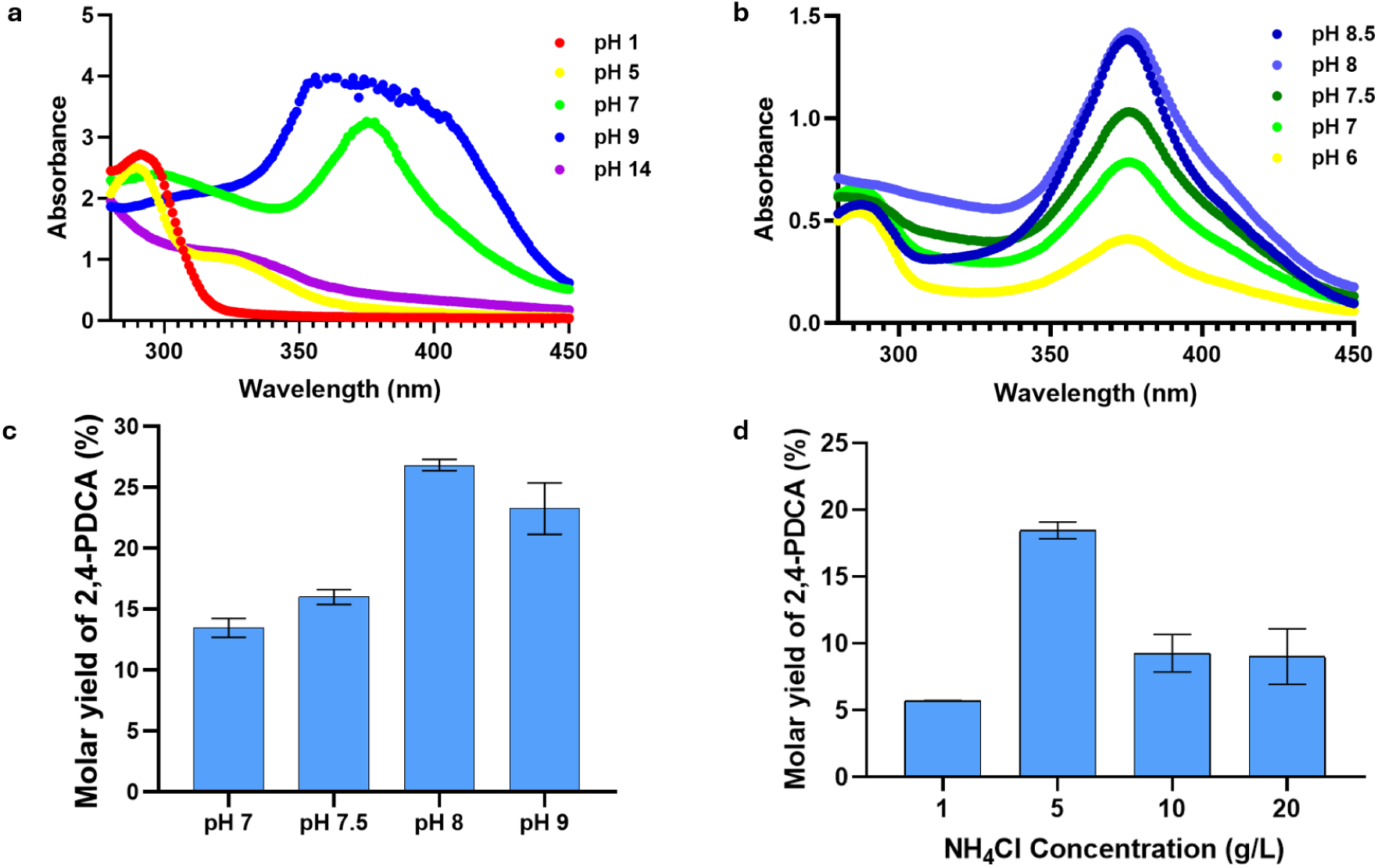
pH and NH_4_Cl concentration optimisation for improving 2,4-PDCA yields. Effect of pH on 4-CHMS detection in **a** and **b** at 380 nm from 132 supernatants of H16 TAPDO *ligAB* resting cell experiments at different pH. **c** molar conversion of TPA to 2,4-PDCA in resting cell experiments (PBS, 2 g/L TPA, 5 g/L NH_4_Cl, OD_600_ 40) at different pH media. **d** the molar yield of 2,4-PDCA from TPA in resting cell conditions (PBS, 2 g/L TPA, pH 7.5 OD_600_ 40) exposed to different concentrations of NH_4_Cl. Molar yields represent means of triplicate values ± SD.

**Figure 4:**
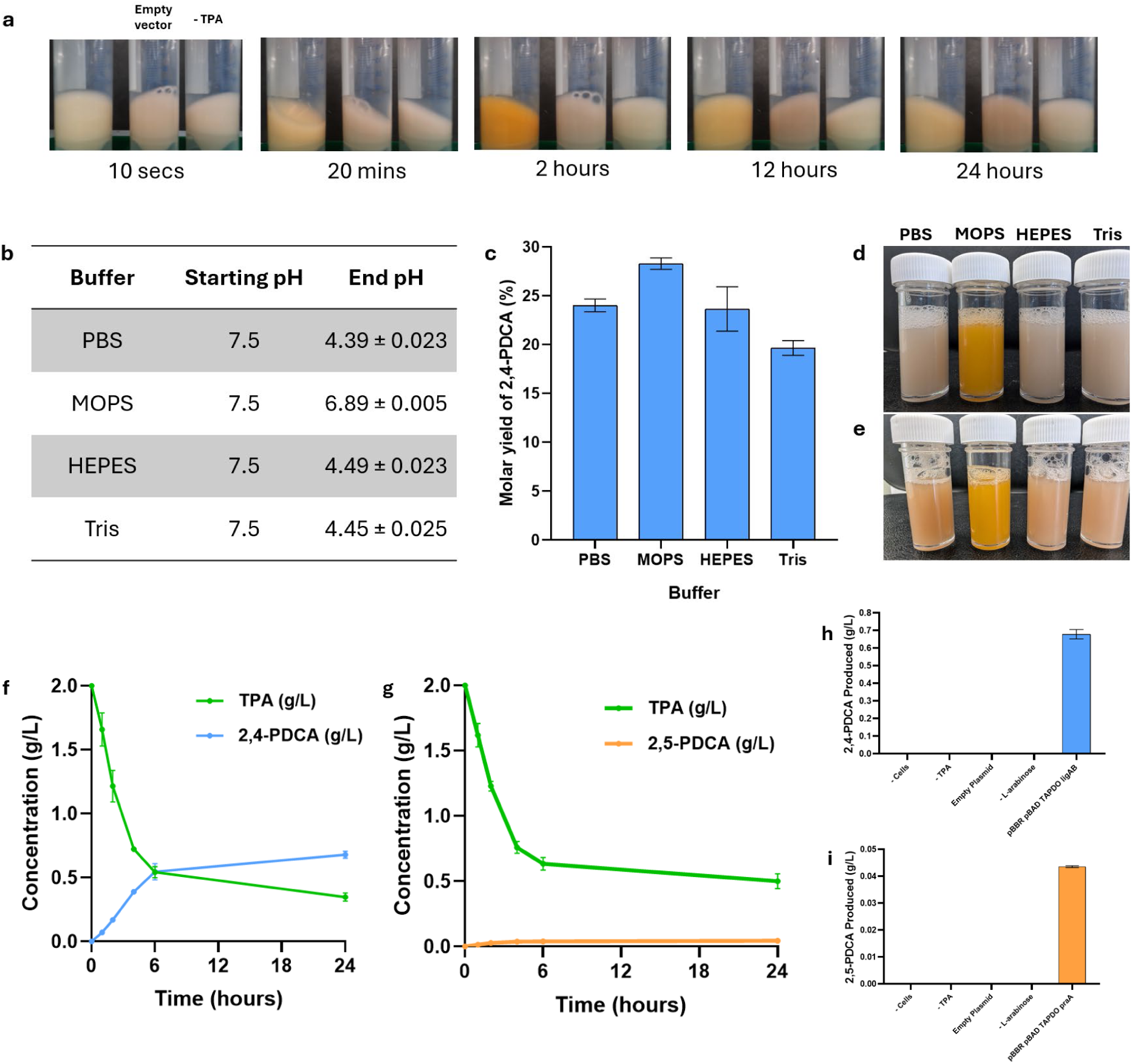
Optimising PDCA production using different buffers. **a** Time course of resting cell reaction with H16 TAPDO *ligAB* (PBS, 2 g/L TPA, 5 g/L NH_4_Cl, pH 7.5, OD_600_ 40) (far left falcon), alongside no TPA (central) and H16 pBBR empty (far right falcon) controls. **b** the start and end pH of resting cell media when using different buffers (PBS, 100 mM MOPS, 25 mM HEPES and 100 mM Tris-base) related to the **c** molar yield of 2,4-PDCA (%) from TPA in these experiments. These media were pictured after 24 hours with media containing PBS, MOPS, HEPES and Tris-base (left to right). **e** These media were then basified using NaOH to pH 9. The optimised production profiles of **f** 2,4-PDCA and **g** 2,5-PDCA was measured from TPA by H16 TAPDO *ligAB* and H16 TAPDO *praA* respectively (100 mM MOPS, 2 g/L TPA, 5 g/L NH_4_Cl, pH 8.0, OD_600_ 40). Concentration of **h** 2,4-PDCA and **i** 2,5- PDCA produced by H16 TAPDO *praA* and H16 TAPDO *ligAB* resting cells compared with control reactions, without cells, without TPA, with empty expression plasmid and without induction. Concentrations are means of triplicate measurements ± SD.

By detecting the presence of CHMS at 380 nm, ideal pH conditions could be determined for producing a greater abundance of its open form between pH 8 and 9 (Figure 3a and b). Increasing the pH of the media, the molar yield of 2,4-PDCA from TPA increased by 13.36 % between resting cell biotransformation undertaken at pH 7 and pH 8 (Figure 3c). To increase cyclisation efficiency higher concentrations of ammonia in the resting cell buffer were also explored, however higher concentrations did not result in higher 2,4-PDCA production (Figure 3d).

The resting cell biotransformations originally undertaken in PBS exhibit rapid formation of a yellow colour that slowly bleaches out over time (Figure 4a). Given PDCAs are not yellow in colour it was hypothesised that this reversion to a colourless state was indicative of final product formation. Resting cell experiments containing PBS drastically changed pH throughout the course of the reactions (from pH 7.5 to 4.4). Thus, different buffers were explored to maintain the abundance of open form CHMS in the basic buffer, whilst the dicarboxylic acid product is formed. It was observed that 25 mM HEPES and 100 mM Tris-base had a similar buffering capacity to PBS (Figure 4 b). Meanwhile, 100mM MOPS had the smallest change in pH throughout the biocatalytic reaction. This improvement was reflected in molar yield of 2,4-PDCA from TPA which increased from 24 % in PBS to 28.3 % in MOPS (Figure 4c). The final colour of the MOPS biotransformation remained yellow after 24 hours. Base (1M NaOH) was added to each of these spent reactions those with weaker buffers exhibited an increased yellow colour (Figure 4d and e), suggesting there is a ‘pool’ of intermediatory carbon existing in the form of closed CHMS that is unavailable for cyclisation to PDCA product due to the more acidic environment. This limits overall conversion of TPA to PDCA product.

Taking into account some of these optimisation strategies, resting cell biotransformations (100 mM MOPS, 2 g/L TPA, 5 g/L NH_4_Cl, pH 8.0, OD_600_ 40) were repeated with H16 TAPDO *ligAB* and H16 TAPDO *praA*. Under these optimised conditions we achieved titres of 0.68 g/L 2,4-PDCA (0.12 g/L·h) and 0.044 g/L 2,5-PDCA (0.012 g/L·h) from TPA over 24 hours (Table 2, Figure 4f and 4g). At no point during these reactions was intermediary metabolite PCA detected. Control experiments did not exhibit PDCA production over the same period (Figure 4i and h).

**Table 2:**
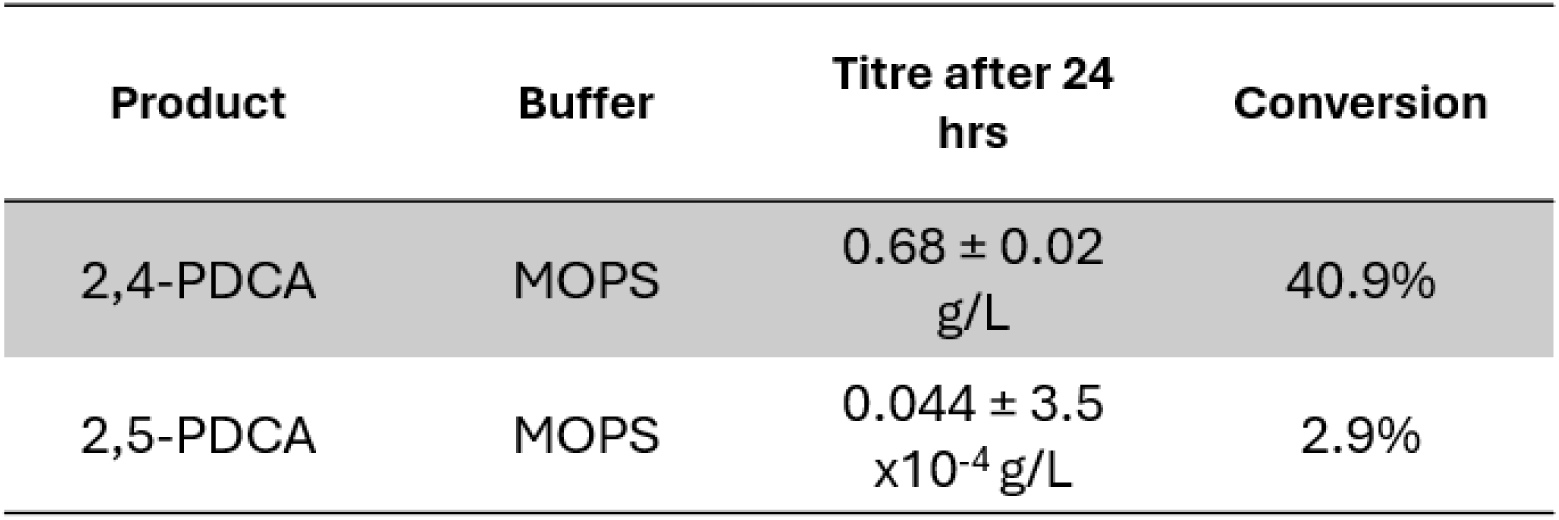
Optimised bioproduction of PDCAs from TPA by H16 resting cells. The yield of PDCAs and molar conversion from 2 g/L TPA by H16 TAPDO *ligAB* and H16 TAPDO *praA* after optimisation (100 mM MOPS, 5 g/L NH_4_Cl, pH 7.5, OD_600_ 40, protein expression = 36 hours). Yields are means of triplicates ± SD. Quantification of metabolites was carried out by HPLC analysis

### Heterotrophic whole cell biocatalysis of TPA to PDCAs

The toxicity exerted by CHMS has resulted in low reported PDCA titres in gram-negative cell chassis, despite advantages to heterotrophic biocatalysis relating to continued protein expression and cofactor renewal. This trend was also observed in this study. H16 TAPDO *ligAB* and H16 TAPDO *praA* were heterotrophically grown and expressed in F-MM (see methods) for 2 hours before being spiked with 2 g/L TPA and 5 g/L NH_4_Cl (pH 7.5, OD_600_ 40). The strains produced 0.14 g/L 2,4-PDCA and 0.015 g/L 2,5-PDCA after 24 hours, with PDCA production mostly taking place in the first 6 hours (Figure 5a and 5b). This premature end to TPA conversion is suggestive of mechanistic inhibition and cell stress via toxic intermediate formation. The cells that converted TPA to 2,4-PDCA in heterotrophic conditions were grown in F-MM for 48 hours alongside cells not exposed to this biocatalytic reaction but that were still undergoing the metabolic burden of induced TAPDO *ligAB* operon. Growth of cells exposed to CHMS was greatly reduced after 48 hours (Figure 5c) which supports the toxic effect of CHMS.

**Figure 5:**
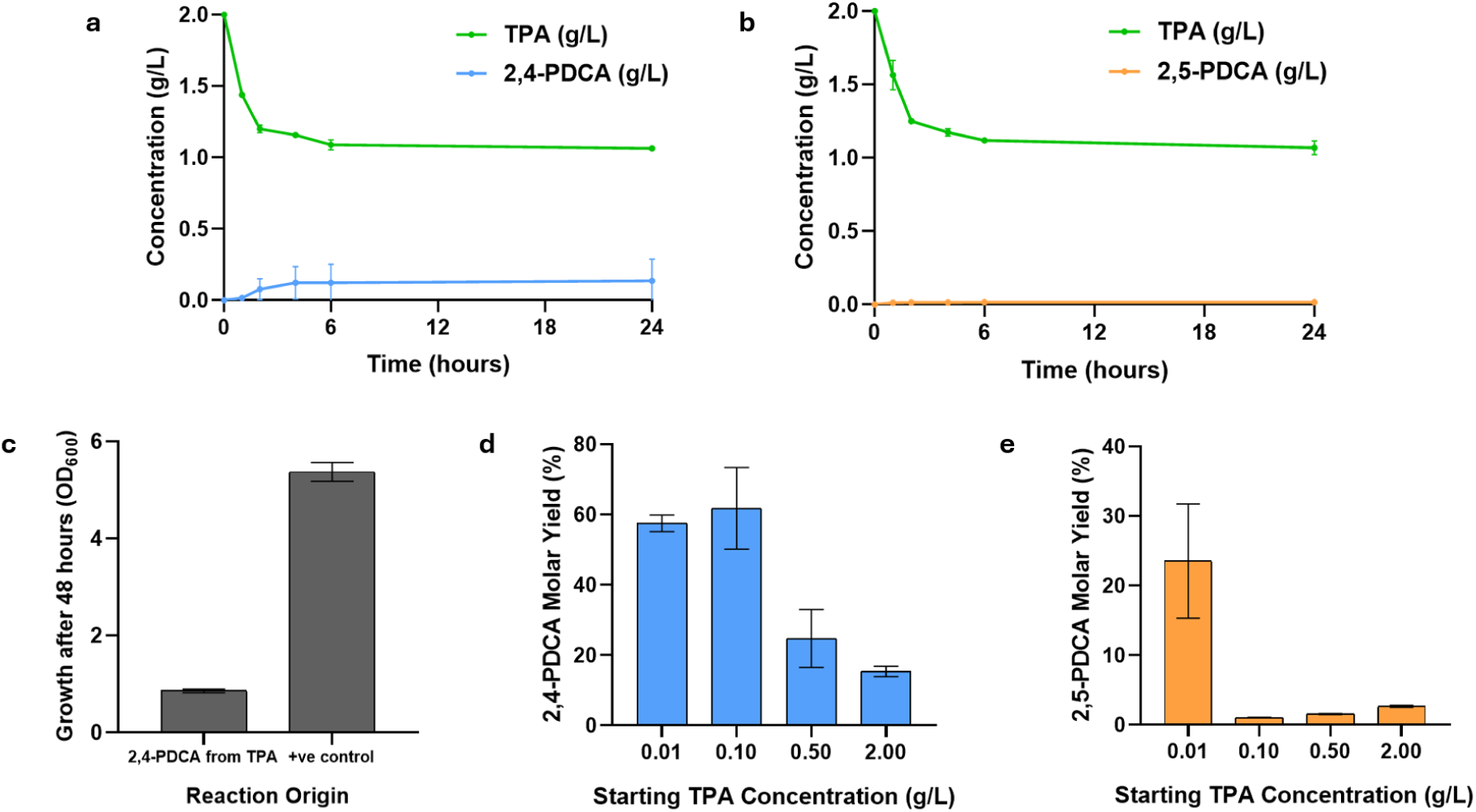
Heterotrophic production of PDCAs by *C. necator* H16. Production of **a** 2,4-PDCA and **b** 2,5-PDCA by heterotrophically growing H16 TAPDO *ligAB* and H16 TAPDO *praA* (F-MM, 2 g/L TPA, additional 5 g/L NH_4_Cl, pH 7.5, OD_600_ 40). **c** OD_600_ of cell cultures inoculated in F-MM from 2,4-PDCA biocatalytic reactions after 48 hours. Heterotrophic production was evaluated for the effect of lowering starting TPA concentrations for improving **d** 2,4-PDCA and **e** 2,5-PDCA molar conversions. All values are average of 3 replicates ± SD.

Attempts to mitigate the inhibitory effects of CHMS accumulation were carried out by reducing the initial concentration of TPA. Figures 5d and 5e highlight greater molar yields of PDCA product when 0.1g/L and 0.01g/L TPA is supplied for (H16 TAPDO *ligAB*) 2,4-PDCA (61.85%) and (H16 TAPDO *praA*) 2,5-PDCA (23.55%) respectively. These higher yields are supportive of the toxicity of CHMS when it accumulates in a larger concentration.

### Influence of *pcaGH* expression

Despite hypothesising the toxicity of CHMS is the main factor in limiting 2,4- and 2,5-PDCA yields, the activity of *pcaGH* is a possible factor for these low yields. *pcaGH* can divert carbon flux away from product formation into the central metabolism (Figure 1). Ideally, a knockout strain of *pcaGH* would be produced but despite several attempts this was not successful. H16 TAPDO *ligAB* and H16 TAPDO *praA* were induced in either LB or F-MM supplemented with TPA. RNA was extracted from these cells during protein expression and during TPA consumption. The rationale of this was to evaluate whether *pcaGH* was upregulated when PCA was present as an intermediate during TPA biocatalysis. The RT-qPCR analysis shows very minimal expression of *pcaGH* both in the absence and presence of TPA. This is signified by the very high comparative expression of *praA* and *ligAB* relative to the background *pcaGH* expression with ∼250,000 fold and ∼3,500-fold higher expression respectively (Figure S1). The relative expression of *praA* (with a 15K RBS translation efficiency), *ligAB* (5K) correlate to their RBS strengths. This high level of transcription relative to *pcaGH* suggests that minimal carbon flux coming from TPA will be diverted into the central metabolism as expression of these production genes outcompete *pcaGH*’s expression and reduce the likelihood of PCA accumulating as a substrate for *pcaGH*.

### PDCA production from TPA under autotrophic conditions

The results discussed above illustrate the successful biocatalysis of plastic monomer, TPA to 2,4-and 2,5-PDCA, albeit at low titres under heterotrophic conditions. Influence of *pcaGH* is also observed to be negligible as its expression is suppressed under heterotrophic conditions which might convert TPA to cell mass. As performed under heterotrophic conditions, where fructose is mainly utilised for cell mass production and maintenance, TPA is diverted towards the PDCAs. In autotrophic cultures, primary metabolism and heterologous protein production will be driven by CO_2_ and H_2_, whereas TPA is diverted towards the PDCAs envisaged towards higher yields of the product. In this way we could demonstrate biocatalysis of TPA in carbon negative mixotrophy.

H16 TAPDO *ligAB* and H16 TAPDO *praA* were grown autotrophically in a bioreactor to OD_600_ 33 and 88 respectively before inducing protein expression for 48 hours. Once spiking with 3 g/L TPA and 5 g/L NH_4_Cl, the vessel was adjusted to pH 7.5 and consumption of TPA was analysed over 24 hours. Following this a further 2 g/L NH_4_Cl was added to the vessel. After 48 hours, 0.22 g/L ± 0.005 2,4-PDCA and 0.12 g/L ± 0.018 2,5-PDCA was produced with maximum productivities of 0.14 g/L·h and 0.05 g/L·h respectively (Figure 6). As previously observed, the majority of PDCA production takes place in the first 6 hours. Interestingly, TPA was entirely consumed in the fed-batch gas fermentation of H16 TAPDO *praA*, however during this time the OD_600_ reduced from 88 to 66 and RT-qPCR of mRNA during this time showed no amplification of *pcaGH*, suggesting carbon flux was not being diverted through the β-ketoadipate pathway. CO_2_ uptake rates between 45-48 mM/h were observed in the cultures which did not change after adding TPA suggesting that the H16 strains use TPA only for PDCA synthesis.

**Figure 6:**
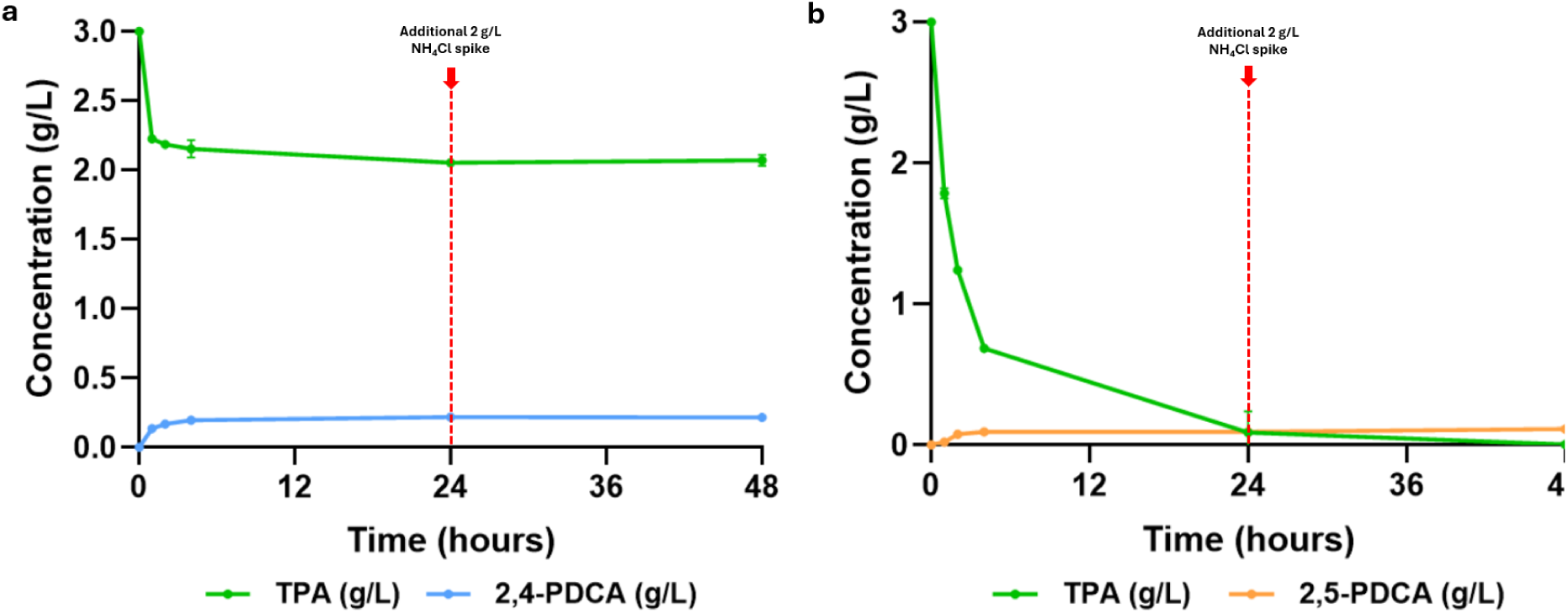
PDCA production from TPA by autotrophically growing *C. necator* H16. In 1 L of a 1.5 L bioreactor **a** H16 TAPDO *ligAB* and **b** H16 TAPDO *praA* were grown on CO_2_, H_2_ and air to OD_600_ 33.6 and 88 respectively before being induced (0.2 % L-arabinose (w/v)) for 48 hours. The vessels were then adjusted to pH 7.5 and spiked with 3 g/L TPA and 5 g/L NH_4_Cl. Samples were taken periodically, and the vessels were spiked again after 24 hours with 2 g/L NH_4_Cl.

### TPA conversion to PDC in *C. necator* H16

Unlike PDCA production, the biosynthetic pathway for PDC production is entirely protein dependent and involves the closed form of CHMS (Figure 7a). PDC is produced from TPA via the overexpression of the TAPDO operon, protocatechuic acid-4,5-dioxygenase *ligAB,* and 4-carboxy-2-hydroxymuconate-6-semialdehyde, *ligC* from *Sphingobium* sp. SYK-6 – producing the strain H16 TAPDO *ligABC.* These genes were expressed under the inducible arabinose promoter in a pBBR expression vector (Figure 7b).

**Figure 7:**
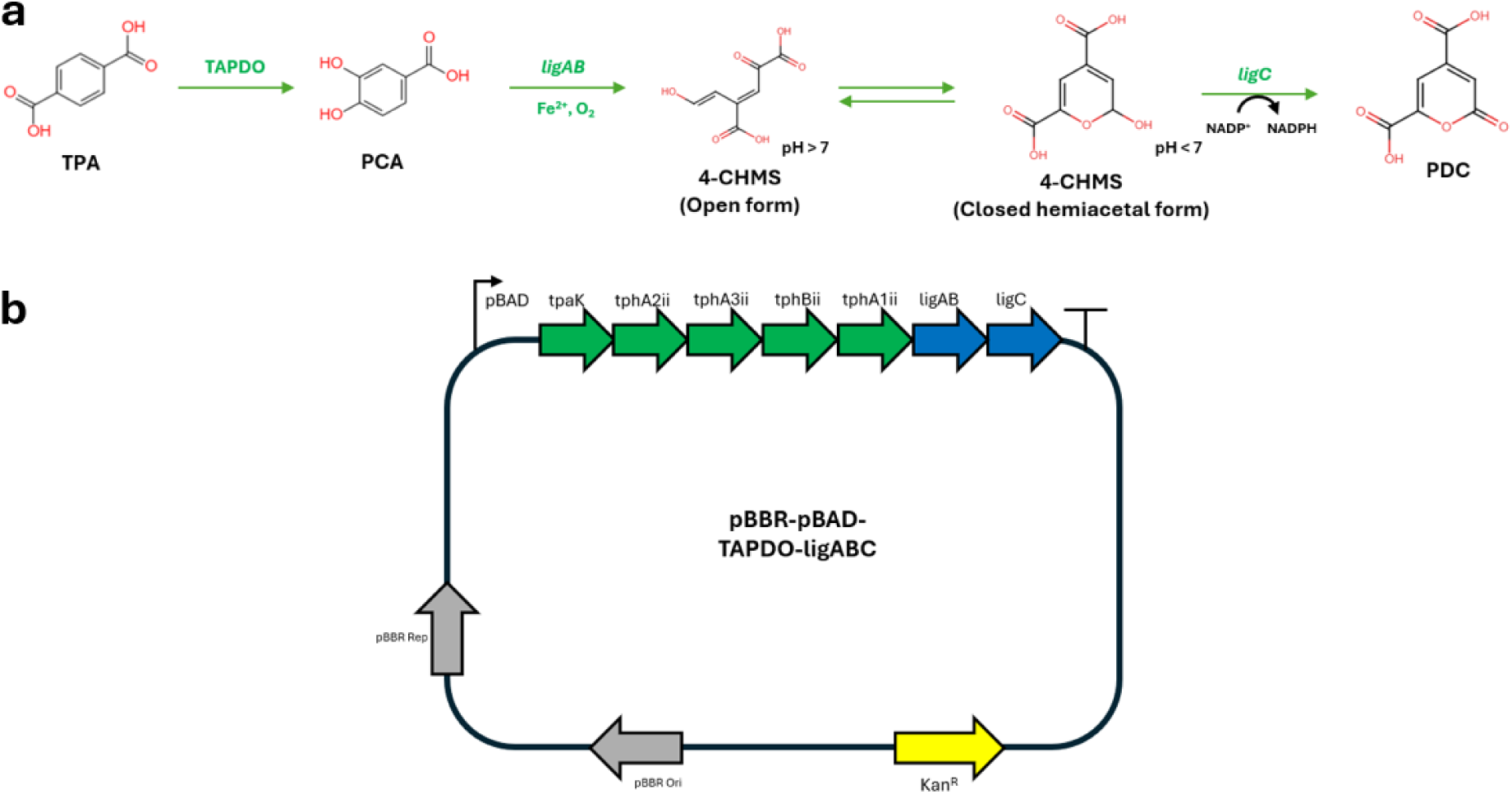
Biosynthetic production of PDC from TPA. **a** Biosynthetic pathway of PDC production from TPA in C. necator H16. The non-native genes expressed (green) are described as is the pH-dependent equilibrium between open and closed forms of 4-CHMS. **b** Schematic illustrating the pBBR expression plasmid containing the TPA converting TAPDO operon (green) and the production genes ligABC (blue) converting PCA to PDC.

Resting cell biotransformations were carried out at pH 5.5 to encourage closed form CHMS to accumulate (Figure 8a). This reaction produced 1.38 g/L PDC from 2 g/L TPA (0.06 g/L·h, 69 % molar yield). The identity of PDC was confirmed by 1H NMR (Figure S2) against previously published spectra and by LC-MS, where an observed mass of PDC (182.99312 m/z) matched the theoretical mass of PDC (182.99297 m/z) (PPM error = 0.81 ppm). Negative control reactions did not produce any PDC after 24 hours (Figure 8b). The success of this reaction was evaluated at more neutral pH to reduce cell stress. These biotransformations produced similar yields between pH 5.5 and 7 (Figure 8c) and broth did not exhibit the yellow colour characteristic of open form CHMS (Figure 8d). This suggests that even in neutral conditions, the pH-dependent equilibrium favours closed form CHMS, and harsher acidic conditions would not be necessary for PDC production.

**Figure 8:**
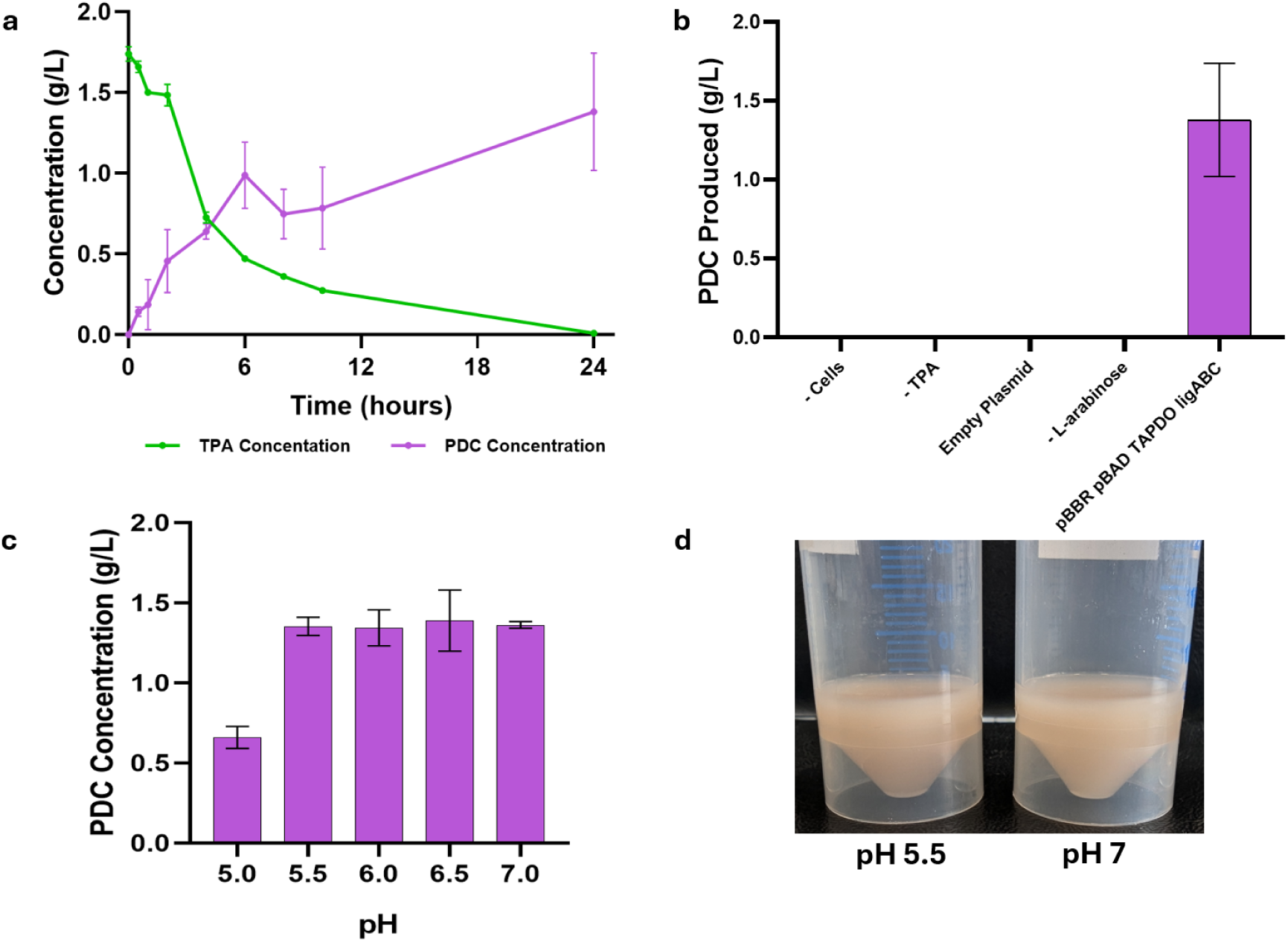
PDC production from TPA by H16 resting cells. **a** Biotransformation of 2 g/L TPA to 1.38 g/L PDC by H16 TAPDO *ligABC* (100 mM MOPS, pH 5.5, OD600 40). **b** Concentration of PDC produced by H16 TAPDO *ligABC* resting cells compared with control reactions, without cells, without TPA, with empty expression plasmid and without induction. **c** The yield of PDC from TPA by H16 TAPDO *ligABC* under resting cell conditions (100 mM MOPS, 2 g/L TPA, OD_600_ 40) at pH 5, 5.5, 6, 6.5 and 7. **d** The colour of the pH 5.5 and 7 resting cell media following these biocatalytic conditions after 24 hours do not exhibit a yellow/orange colour characteristic of open form CHMS. Concentrations are represented as means of triplicates ± SD.

### Enhancing PDC production from TPA in heterotrophically growing *C. necator* H16

PDC production from heterotrophically growing H16 was expected to be successful due to a reduction in pathway toxicity and continued protein expression. H16 TAPDO *ligABC* growing in F-MM (pH 6.5, OD_600_ 40) was spiked with 2 g/L TPA. PDC was rapidly produced at 0.35 g/L·h, with a final titre of 1.93 ± 0.05 g/L after 24 hours, constituting a 96.5 % molar yield of PDC from TPA (Figure 9a). Lower starting concentrations of TPA led to equally high PDC production, suggesting absence of toxic intermediates in the PDC production pathway (Figure 9b).

**Figure 9:**
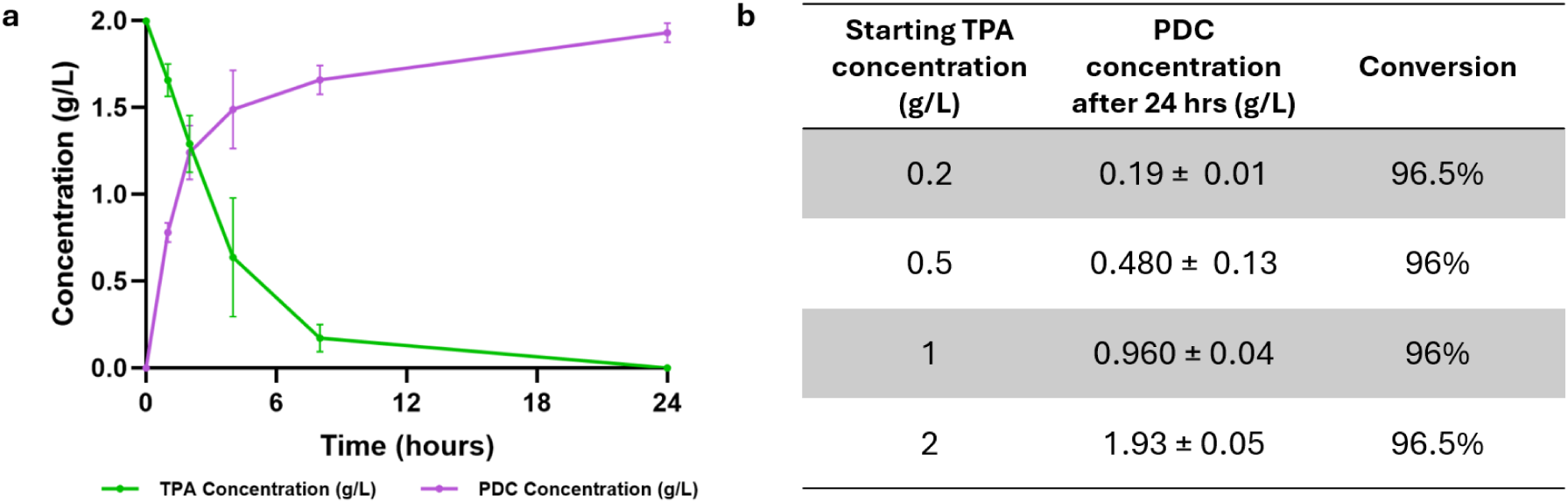
PDC production from TPA by heterotrophically growing H16. **a** 1.93 g/L PDC produced from TPA in H16 TAPDO *ligABC* cells growing in F-MM (0.2% (w/v) L-arabinose, pH 6.5) for 2 hours before being spiked with 2 g/L TPA. **b** Conversion of TPA to PDC with different starting concentration of TPA. Mean concentration of PDC after 24 hours from 0.2, 0.5, 1 and 2 g/L TPA by heterotrophically growing H16 TAPDO *ligABC* (F-MM, 0.2% (w/v) L-arabinose, pH 6.5) ± SD.

### Continuous mixotrophic gas fermentation for TPA conversion to PDC

The batch and fed-batch bioproduction of PDC from vanillic acid and glucose has been successfully explored to high titres, but these cell factories release biogenic CO_2_ during metabolism that reduces carbon yields and overall sustainability. A continuous fermentation bioprocess has the potential for greater productivity and economic feasibility. Establishing PDC production in a carbon negative continuous fermentation has not yet been explored. *C. necator* H16 is able to grow in mixotrophy on CO_2_ and H_2_ as well as other organic carbon sources. This study’s main aim is to use CO_2_/H_2_ for cell mass and protein production, TPA for target product formation with equimolar conversion yields, uncoupling cell growth and product formation. If this can be achieved in a continuous process, that would be beneficial for a techno-economically feasible bioprocess.

H16 TAPDO *ligABC* was grown autotrophically in a gas fermentation system as described in the methods. The culture was induced by adding L-arabinose at OD_600_ 66 and expressed for 36 hours before starting to feed the bioreactor with minimal media supplemented with 21.5 g/L TPA and 0.2 % (w/v) L-arabinose. The dilution rate of the vessel was 0.02 h^-1^ which allowed the culture to reach steady-state at OD_600_ ∼165 (Figure 10). Given TPA was previously consumed at a maximum rate of 0.35 g/L·h (Figure 9a), a feed rate of ∼0.43 g/L·h was chosen to minimise TPA accumulation, that could increase *pcaGH* expression and divert carbon flux into the central metabolism. Indeed, throughout the fermentation, an excess of 0.006 g/L TPA was detected and no accumulation of PCA was observed. CO_2_ uptake remained constant (50 mM/h) before and during exposure to TPA. This suggests that the carbon flux from TPA remained unchanged and *pcaGH* expression did not increase when the PDC production pathway was active. This was confirmed by RT-qPCR, where amplification of *pcaGH* was not observed. Hydrogen uptake rate of 273 mM/h and oxygen uptake rate of 125 mM/h was observed during the steady state (Figure S3).

**Figure 10:**
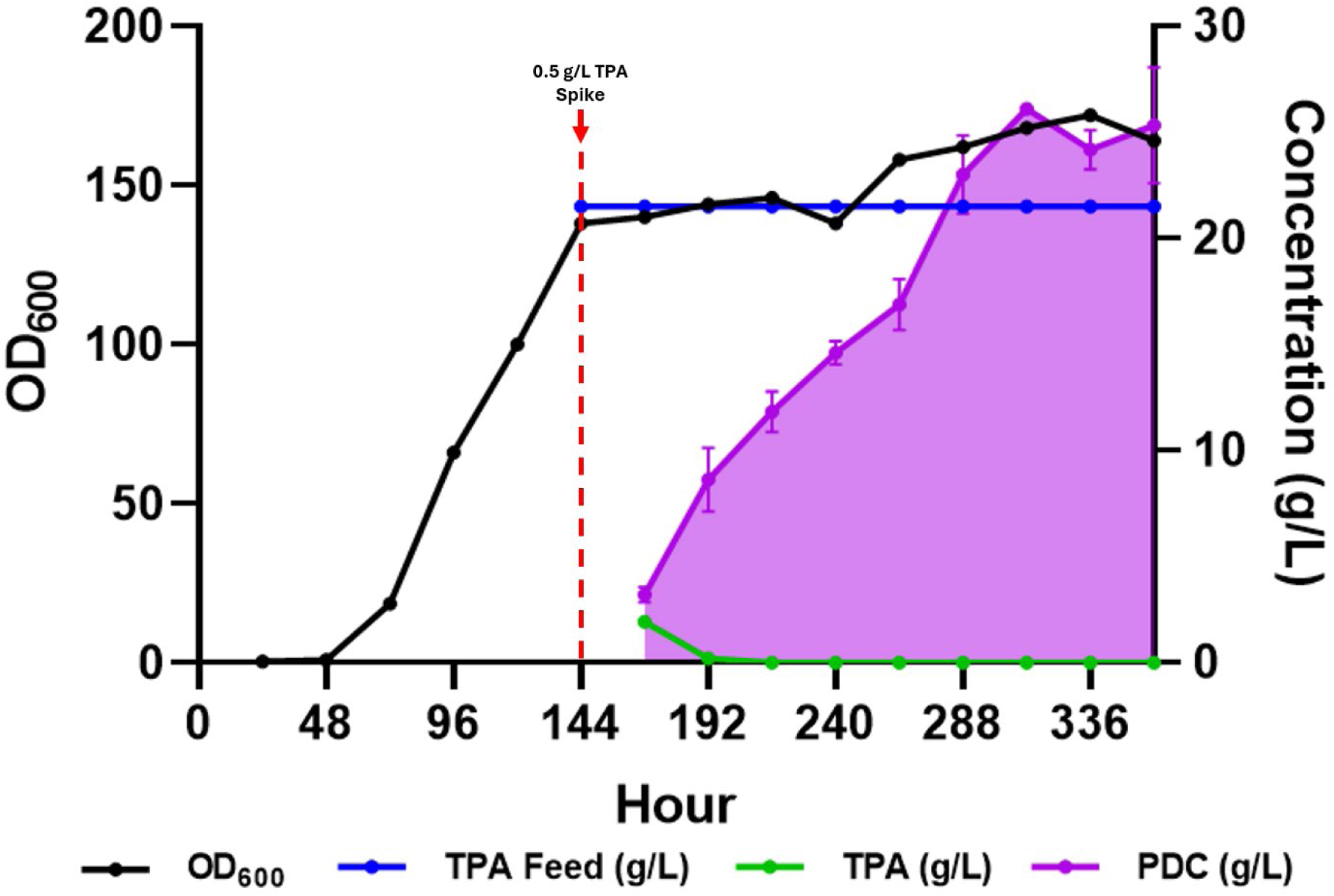
Continuous mixotrophic PDC production from TPA by CO_2_-driven H16. Growth (OD_600_) of H16 TAPDO *ligABC* (black line) measured at each time point. The red line (hour 144) signifies the beginning of continuous fermentation at a vessel dilution rate of 0.02 h-^1^ by feeding media supplemented with 21.5 g/L TPA (0.43 g/L·h) over 24 hours (blue line). Unreacted TPA is represented in green whilst PDC production is highlighted by the shaded area (purple), and are means of triplicate measurements ± SD.

When fed with 0.43 g/L·h TPA, PDC rapidly accumulated in the first 5 days of continuous fermentation before stabilising at 24.65 ± 1.43 g/L PDC (0.49 ± 0.03 g/L·h) in the final 4 days (Figure 10). With an actual TPA consumption rate of 0.42 g/L·h, the maximum theoretical steady state concentration of PDC produced would be 23.48 g/L (0.47 g/L·h). This continuous mixotrophic fermentation achieved ∼100% molar conversion of TPA to PDC in a carbon negative bioprocess, with the highest reported yield of PDC under continuous autotrophic fermentation and highest reported titre and productivity metrics using TPA as feedstock.

## Discussion

This study demonstrates the first carbon-negative bioprocess for converting PET monomer, terephthalic acid into value-added dicarboxylic acids using mixotrophic gas fermentation. By leveraging the metabolic versatility of *C. necator* H16, we achieved complete conversion of TPA to PDC with techno-economically feasible metrics while simultaneously fixing CO₂, addressing both plastic waste valorisation and carbon capture in a single integrated process.

The key innovation in our approach lies in uncoupling biomass formation from product synthesis through mixotrophic cultivation. Traditional heterotrophic bioconversions of aromatic compounds suffer from carbon loss to biomass and CO₂, limiting theoretical yields to ∼60-70%[20]. Our system overcomes this limitation by utilizing CO₂ and H₂ for cell growth via the Calvin cycle and heterologous protein synthesis, while channelling TPA exclusively toward product formation. This metabolic division of labour enabled near-quantitative conversion of TPA to PDC (∼100% molar yield), exceeding previously reported yields and productivities from sugar-based as well as using TPA as feedstocks [12, 21]. The study by Otsuka et al based on using lignin derived vanillin did achieve 99% molar yields of PDC with high titres (99 g/L) and productivities, where glucose was used to support primary metabolism[16]. For high cell densities using sugar-based feedstocks there is significant amount of CO_2_ that is being released within this process.

Kang et al [17] demonstrated a chemo-hybrid process for converting waste PET to PDC. In this study, chemo-catalytic conversion of PET to pure TPA was demonstrated which is then used as a feedstock for two *E. coli* strains, one carrying the TPA to PCA module, and the second one carrying the PDC module. The two-strain strategy demonstrated a proof-of-concept conversion of TPA to PDC achieving 99% conversion yields, ∼0.3 g/L·h productivities albeit low titres. In another study *Comamonas testosteroni* was engineered to produce PDC from TPA and glycolic acid[21]. This study demonstrated productivities of 0.27 g/L·h where the overall efficiency reported is about 100%.

The continuous fermentation data from the current work provide compelling evidence for the decoupling strategy between growth and product formation. CO₂ uptake rates remained constant at about 50 mM/h before and during TPA feeding (Figure S3), indicating that autotrophic metabolism sustained cellular maintenance while TPA was funnelled entirely to PDC. The absence of *pcaGH* expression during steady-state production further confirms that carbon flux through the β-ketoadipate pathway was effectively blocked, preventing TPA catabolism for growth. Although this work indicated that *pcaGH* knockout is not essential, this would have been beneficial for the PDCAs production as TPA conversion only accounted to maximum of 22%.

The stark difference in production efficiency between PDC (100% yield) and PDCAs (22% and 4% for 2,4- and 2,5-PDCA, respectively) reveals fundamental constraints in engineering aromatic metabolism. Both pathways share the toxic intermediate 4- or 5-carboxy-2-hydroxymuconate-semialdehyde (CHMS), but their divergent pH requirements create opposing optimization challenges. PDC formation requires the closed form of CHMS (favoured at pH <7), enabling enzymatic conversion by *ligC*. In contrast, PDCA synthesis depends on spontaneous cyclization of open-form CHMS with ammonia, requiring pH >8. In previous studies [12, 18], 2,4- and 2,5-PDCAs were produced from TPA and glucose respectively. In both cases after optimising conditions such as pH and ammonia concentrations the maximum titres obtained were 0.5 g/L 2,4-PDCA and 1.4 g/L 2,5-PDCA from glucose. This suggests certain metabolic limitations of either the availability of free ammonia within the cells to perform the cyclisation into PDCAs or even the conversion of CHMS to other products such as picolinate. A recent study has shown that the 5-CHMS is decarboxylated to picolinate and is the preferred route in *E. coli*[13]. In this work, an enzyme-based ammonia incorporation has enhanced the 2,5-PDCA titres to 10 g/L from glucose and pyruvate.

Our optimization studies revealed that even with improved buffering (100 mM MOPS) and elevated pH, PDCA yields remained limited. The accumulation of “trapped” CHMS intermediates evidenced by the yellow coloration that intensified upon basification suggests that the spontaneous cyclization rate is intrinsically slower than enzymatic PDC formation. This mechanistic bottleneck, combined with CHMS toxicity at alkaline pH, fundamentally limits PDCA production in *C. necator*. Future strategies might explore protein engineering of ring-cleavage dioxygenases to reduce CHMS accumulation or screening for enzymes that catalyse PDCA formation rather than relying on spontaneous chemistry.

The continuous fermentation achieving 24.5 g/L PDC at 0.47 g/L·h productivity represents the highest reported PDC titre from TPA and the first demonstration of continuous autotrophic based PDC production. Here we also have to imply that the PDC production is limited by TPA feed concentration, future experiments will need to identify the maximum TPA feed concentration which could allow to reach high titres and productivities of PDC in the continuous autotrophic cultivations.

More significantly, this process exemplifies a new paradigm for carbon-negative biomanufacturing. By maintaining the culture at OD₆₀₀ ∼165 through autotrophic growth while continuously converting TPA to product, we achieved several critical advantages. These include a stable culture for more than 4 days without productivity decline and no PCA accumulation or TPA diversion towards cell mass. This high cell density, combined with efficient enzyme expression, enabled equimolar TPA conversion to PDC exceeding previously reported metrics using TPA[21]. Moreover, the *C. necator* H16 background strain also produces PHB as its natural product. In this scenario, the product sinks are ideally cofactor balanced as 1 mol of NADPH is produced by *ligC* in the PDC pathway which could be recycled via the PHB pathway. Moreover, our study shows that in *C. necator*, TPA can be used under heterotrophic conditions with an organic carbon source such as fructose and this can also replicate similar observation where fructose was mainly used for biomass formation and protein production, TPA diverted towards PDC with more than 95% conversion efficiency (Figure 9b).

Our results establish mixotrophic gas fermentation as a viable strategy for integrating waste valorisation with carbon capture. The process simultaneously addresses two pressing environmental challenges: plastic pollution and atmospheric CO₂ accumulation. With global PET production exceeding 70 million tons annually and enzymatic recycling gaining momentum, TPA availability as a feedstock will increase substantially. Converting this waste stream to PDC, a precursor for biodegradable polymers, pharmaceuticals, and agrochemicals creates economic incentives for plastic collection and recycling. Techno-economic feasibility of enzymatic depolymerisation of PET has been showcased where if this is combined with an efficient conversion of TPA to a biopolymer would further enhance the economic feasibility and environmental impact[22].

The techno-economic viability of this process is enhanced by several factors: (1) TPA requires no pretreatment beyond PET hydrolysis, unlike lignocellulosic feedstocks; (2) continuous operation minimizes fermentation infrastructure; (3) product recovery is simplified by PDC’s high water solubility and stability; (4) the process generates no waste streams, with all carbon directed to product or biomass. The metabolic architecture developed here which is a continuous autotrophic growth supporting heterologous biocatalysis, could be extended to other complex substrates and products which will enable true carbon negative sustainable bioprocess.

## Conclusion

This work demonstrates that combining gas fermentation with plastic monomer valorisation creates a carbon-negative route to valuable chemicals. The complete conversion of TPA to PDC via mixotrophic cultivation of *C. necator* H16 validates the concept of decoupling growth from production in gas-fermenting chassis. Limitations within the PDCA production pathways needs further research in reducing the toxicity of open form CHMS and the dependence on ammonia for cyclisation. While PDCA production remains limited by pathway-specific constraints, the PDC production pathway achieved near-complete TPA conversion. As the circular economy transitions from concept to implementation, such integrated bioprocesses that transform multiple waste streams into products will become increasingly vital. Our findings provide both a specific solution for TPA valorisation and a general strategy for designing carbon-negative biomanufacturing processes that align economic incentives with environmental sustainability.

## Methods

### Chemicals, bacterial stains, plasmids, culture conditions

All chemicals were purchased from Thermo Scientific, Sigma-Aldrich or Melford unless otherwise stated. *E. coli* strains DH5α and S17-1 were used for plasmid propagation and conjugative plasmid transfer respectively. *E. coli* strains were cultivated at 37 °C in LB medium supplemented with 50 µg/ml kanamycin when carrying a plasmid. *C. necator* H16 strains were grown in LB medium or minimal media (F-MM) [23] at 30 °C with 10 µg/ml gentamycin. 20 g/L fructose was used as carbon source unless otherwise stated. Bacterial strains and plasmids used in this work are stated in Table S1.

### Autotrophic media composition

The minimal media used for autotrophic growth of C. necator H16 consisted of 3.4 g/L Na_3_P_3_O_9_, 1.5 g/L K_2_SO_4_, 1.5 g/L NH_4_Cl, 0.5 g/L MgSO_4_.7H_2_O, 0.01 g/L CaCl_2_, 0.05 g/L Fe (III) NH_4_-Citrate and 5 ml/L SL6 [5].

### Construction of expression plasmids and gene expression

Expression of the PDCA and PDC producing pathways was carried out using the vector pBBR1-USERcassette1. The vector was linearised using XbaI and then Nt. BbvCI before being assembled with the relevant genes (*araC*/P_BAD_, *tpaK, tphA2II, tphA3II, tphBII, tphA1II, praA, ligAB, ligC*). Plasmids were conjugated into *C. necator* H16 using *E. coli* S17-1 as donor strain. Donor and recipient strains were allowed to mate for 6 hours before transconjugants were selected on fructose-MM agar plates supplemented with 10 µg/ml gentamycin and 300 µg/ml kanamycin.

Apart from araC/P_BAD_, all genes were codon optimised for *C. necator* H16 and synthesised (GenScript UK Limited) into pCU57 plasmids (Table S2). All primers were synthesised by IDT (Table S3). Relevant genes were amplified using Phusion U Green Hot Start DNA Polymerase (Thermo Scientific) and purified by gel extraction. Assembly of expression plasmids was carried out by USER Cloning (New England Biolabs). Following cloning, DNA constructs were confirmed by both PCR and sanger sequencing (Eurofins Genomics).

Following successful assembly and conjugation, protein expression was confirmed by reverse transcriptase quantitative PCR (RT-qPCR) of mRNA. After strains were subject to protein expression in either LB medium or fructose-MM (supplemented with TPA), RNA was extracted and purified using Monarch Total RNA Miniprep Kit (New England Biolabs). RT-qPCR was carried out using Luna® Universal One-step RT-qPCR kit (NEB) according to manufacturer’s protocols. Reactions used gene specific primers and an internal control of 16s rRNA that was then visualised using an Applied Biosystems StepOnePlus® instrument.

### PDCA and PDC production in *C. necator* H16 resting cells

H16 carrying pBBR pBAD TAPDO *praA*, pBBR pBAD TAPDO *ligAB* or H16 pBBR pBAD TAPDO *ligABC* were grown in LB medium supplemented with 0.2% (w/v) L-arabinose and necessary antibiotics. After 16-48 hours of incubation shaking at 30 °C cell cultures were pelleted and washed twice in 100 mM MOPS (unless otherwise stated) before being inoculated in resting cell buffer to a starting OD_600_ of 40. Regarding 2,4-PDCA and 2,5-PDCA production, the optimum resting cell buffer for these strains consisted of: 100 mM MOPS, 2 g/L TPA, 5 g/L NH_4_Cl at a pH of 7.5 unless otherwise stated. When producing PDC under resting cell conditions, the inoculated buffer contained: 100 mM MOPS, 2 g/L TPA at a pH of 5.5 unless otherwise stated. Experiments were performed in triplicates and samples were taken at various time points and stored at -20 °C before downstream analysis. Negative controls without the addition of TPA substrate or the use of H16 pBBR empty vector were used for the validity of this experiment.

### PDCA and PDC production in *C. necator* H16 heterotrophically growing cells

Precultures of H16 carrying pBBR pBAD TAPDO *praA*, pBBR pBAD TAPDO *ligAB* or H16 pBBR pBAD TAPDO *ligABC* were grown and washed as described above. Cells were then inoculated into F-MM (20 g/L fructose, 0.2 % (w/v) L-arabinose, OD_600_ 40, pH 7.5) for two hours, stimulating heterotrophic growth on fructose. After two hours, cultures were spiked with 2 g/L TPA and an additional 5 g/L NH_4_Cl.

### PDCA and PDC production in *C. necator* H16 autotrophically growing cells

The autotrophic fermentations were performed in a Applikon ez2-Control bioreactor system (Getinge) and the set-up is similar to that was described previously [5]. A double jacked 1.5 litre vessel with a working volume of 1.0 litre was used. Off-gas analysis was performed by a Raman based gas analyser (Atmospheric Inc.). The peristaltic pumps and different width tubing was calibrated to relate pump RPM to ml/hour of media. The culture temperature was maintained at 30 °C, the pH was maintained at 6.8 using a pH probe. The agitation speed was initially set to 400 rpm with CO_2_ (30 ml/min), H_2_ (780 ml/min) and air (190 ml/min) fed into the vessel. As strains began to grow, agitation and airflow then became coupled to dissolved oxygen, this allowed for the support of growth at higher ODs. When necessary, protein expression was induced with the addition of sterile 0.2% (w/v) L-arabinose. When operating under batch mode, the initiation of metabolic pathways was started by the addition of starting substrate into the vessel. Time-course samples prior and during these reactions were taken for analysis. The initiation of continuous mixotrophic fermentation was carried out by the connection of feed media and waste to the vessel. The vessel was spiked with the desired substrate, and the feed media contained the substrate at the same concentration (g/L·h) along with L-arabinose (0.2 % w/v) to support continued protein expression. The feed-media was fed into the vessel at a specified dilution rate via a peristaltic pump. The waste was then automatically drawn from the fermenter vessel to maintain the working volume at 1 L.

A 5 ml overnight culture of either H16 pBBR pBAD TAPDO praA, ligAB, or ligABC were grown and inoculated into 50 ml of LB in a baffled flask and grown overnight. This 50 ml culture was used unwashed to inoculate the bioreactor containing 1 L fermenter minimal media. Cells were grown to OD_600_ ∼ 60-100 and left to express for 36 hours. At this stage continuous fermentation could begin, the vessel was spiked with TPA substrate (to the same hourly concentration as what was in the feed) and feed media (minimal media, 21.5 g/L TPA (may vary), 0.2 % (w/v) L-arabinose, additional 5 g/L NH_4_Cl when necessary) was fed into the vessel at a dilution rate of 0.02 h^-1^ unless otherwise specified. A bleed line from the vessel was attached leading to a waste bottle, allowing for the volume to be maintained at 1L. For PDCA production the pH was adjusted to 7.5 before feeding, pH 6.5 was used for PDC production. During continuous fermentation, steady state was reached after ∼3 days, with product accumulation monitored throughout the fermentation.

Fed-batch PDCA production took place in the same way, except after 36 hours of protein expression, the vessel was directly spiked with 3 g/L TPA (from a 100 g/L stock) and 5 g/L NH_4_Cl (from a 200 g/L stock). PDCA production was then monitored over the following 24 hours. After 24 hours, the vessel was spiked with an additional 2 g/L NH_4_Cl and the vessel was sampled for a further 24 hours before stopping the run.

### Metabolite analysis

Growth of cell cultures were monitored by their optical density at a wavelength of 600 nm (OD_600_) using a spectrophotometer. The concentrations of TPA, PCA, 2,4- and 2,5-PDCA were determined on a Thermo Vanquish HPLC-UV-IR instrument equipped with a InfinityLab Poroshell 120 Aq-C18 (Aligent) (4.6 mm x 150 mm, 2.7-micron) column. A gradient method with a flow rate of 1 mL/min was used with mobile phase A (0.025% H_3_PO_4_) and mobile phase B (100% acetonitrile). Phase B increased from minutes 0 to 13.8 from 0 to 69%. This percentage then increased to 100% over the next 30 seconds and remaining at 100% for 3 minutes before decreasing to 0 % in 30 seconds and remaining at 0% for the remaining 6.5 minutes. Key metabolites were detected at 210, 260 and 280 nm.

Concentrations of PDC were determined by ^1^H NMR. ^1^H NMR was recorded on a JEOL ECS400FT Delta spectrometer (at 399.78 MHz). Peaks were referenced against a D_2_O internal standard containing 10 mM trimethylsilylpropanoic acid. JEOL Delta (v6.3) software was used for interpretation of spectra and quantification of PDC.

Intermediates 4- and 5-CHMS were detected by a Spark Multimode Microplate reader (Tecan) using samples in a Greiner 96 well UV transparent microplate. Absorbance scans of samples were undertaken with CHMS exhibiting a UV-Vis spectrum at a λmax of 380 nm.

Mass spectroscopy analysis was conducted to further confirm the identity of metabolites PDC, 2,4- and 2,5-PDCA. Profiling was conducted on a Vanquish UHPLC system with a Thermo IDK Tribird high resolution mass analyser (Waltham, MA, USA). Sample separation was achieved using a high-strength silica column (1.8 µm x 2.1 mm x 150 mm) operating at 35 °C. With a 5 µL injection volume and 250 µl/min flow rate, buffer A (95 % acetonitrile) and buffer B (0.1 % ammonium hydroxide) were used in a gradient method, with a 1.5-minute hold of 95 % A/5 % B followed by a decrease to 5 % A/95 % B at 11.5 minutes. After, a washing cycle hold for 5 minutes, at 16.1 minutes proportions return to 95 % A/5 % B. Data acquisition settings were as follows for MS1 acquisition: Orbitrap analyser, mass resolution 30 K, quadrupole isolation, scan range 100-1000 m/z, RF lens 30 %, Automatic gain control 95 %, injection time 50 ms. Isolation Mode (Quadrupole): 1.5 m/z, isolation off (off), HCD stepped energy (10, 35, 60), detector type (orbitrap, 30 K mass resolution), AGC target 50% and maximum injection time 50 ms.

## Supplementary information

### Supplementary Figures

**Figure S1:**
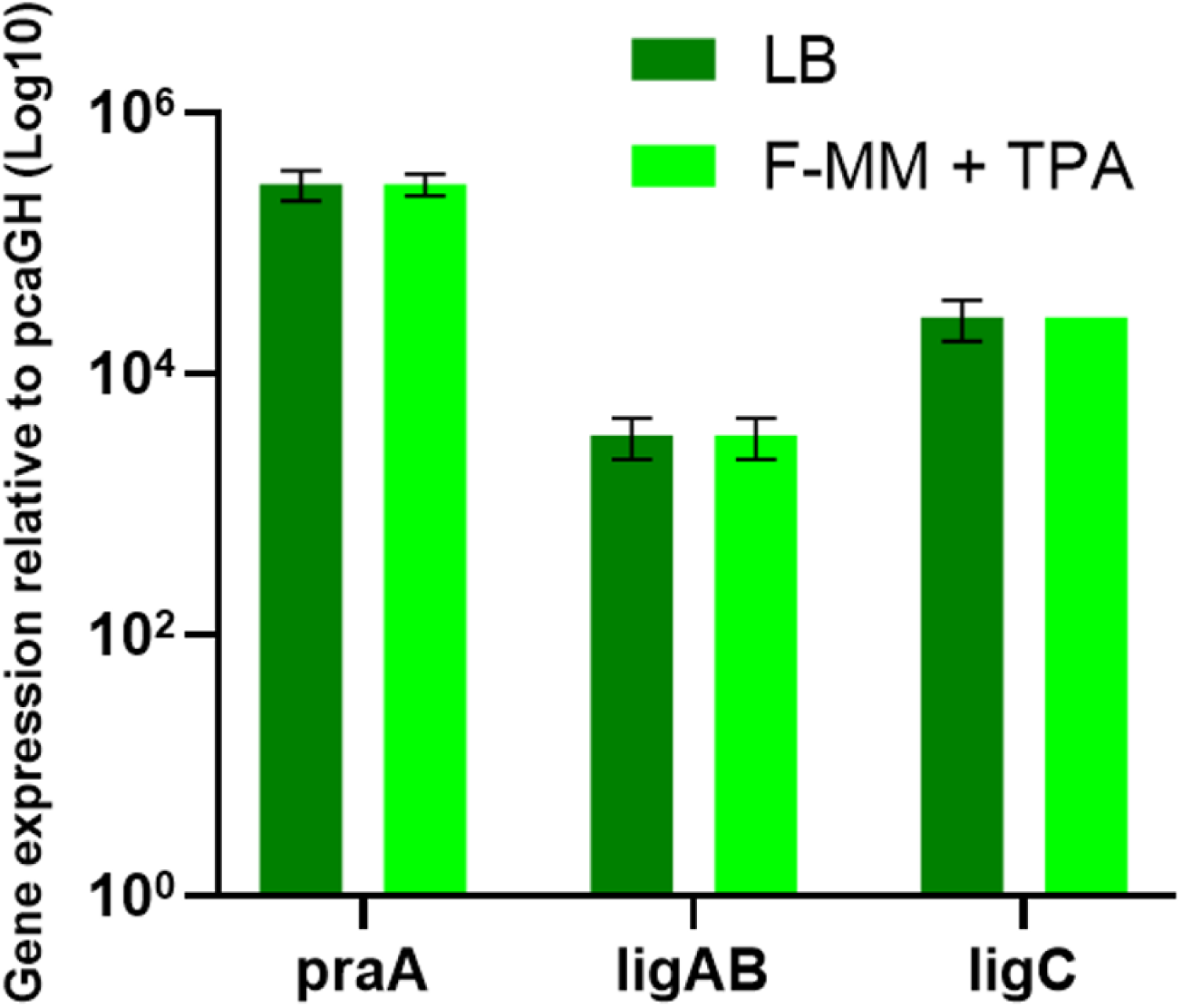
RT-qPCR analysis of production genes in 2,4-PDCA, 2,5- PDCA and PDC biosynthetic pathways relative *pcaGH* expression. The mRNA expression level of H16 TAPDO strains expressing production genes *praA*, *ligAB* and *ligC* was normalised against 16s rRNA gene expression and then against *pcaGH* expression. This was done using expressing cells in LB media and in F-MM during the consumption of TPA to see any influence of TPA on *pcaGH* expression. The results are means of triplicates ± SD.

**Figure S2:**
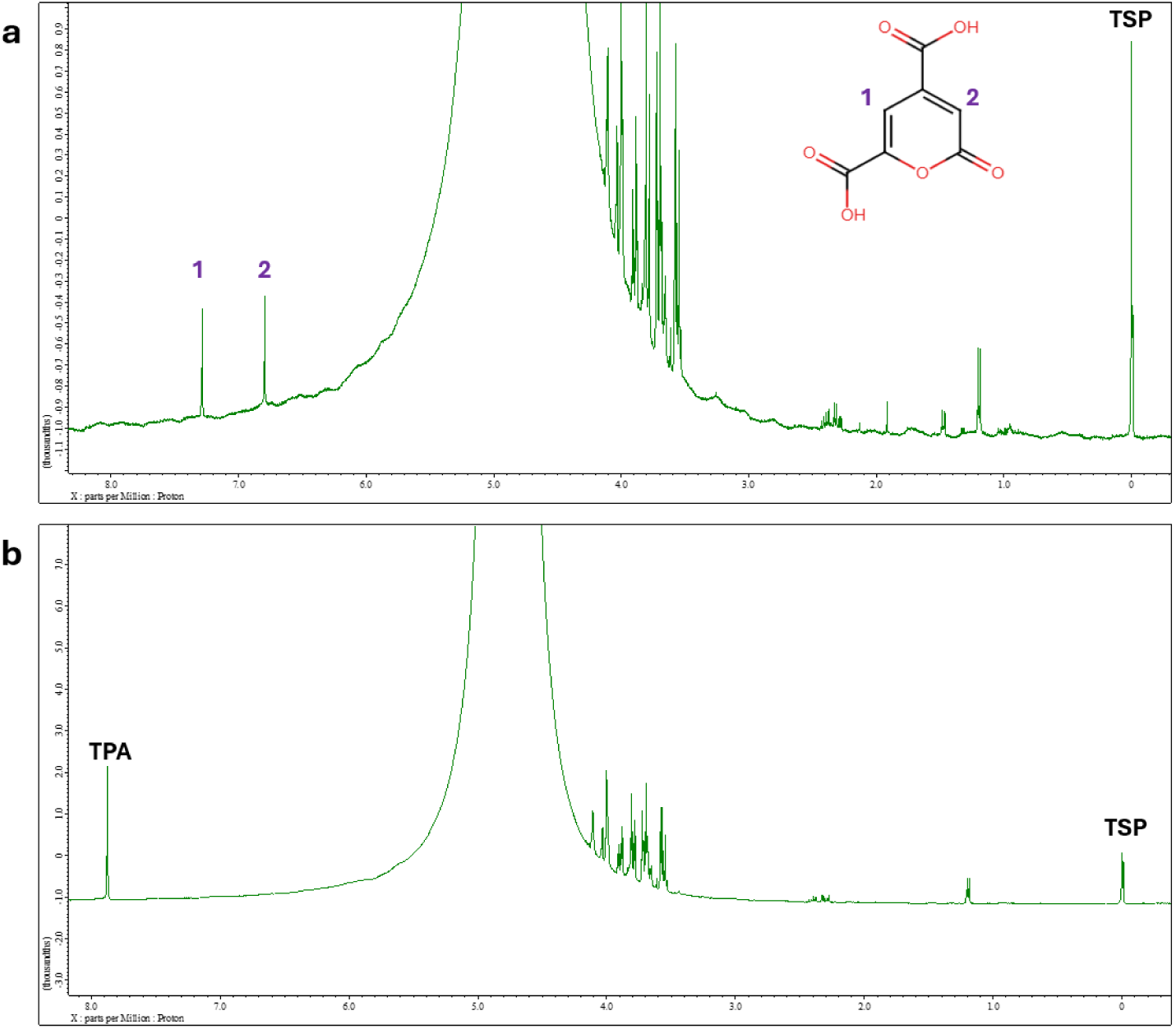
^1^H NMR spectra of PDC. Samples from TPA biotransformations inoculated with **a** H16 TAPDO *ligABC* and **b** H16 pBBR empty were collected after 24 hours and subject to NMR demonstrating authentic PDC production.

**Figure S3:**
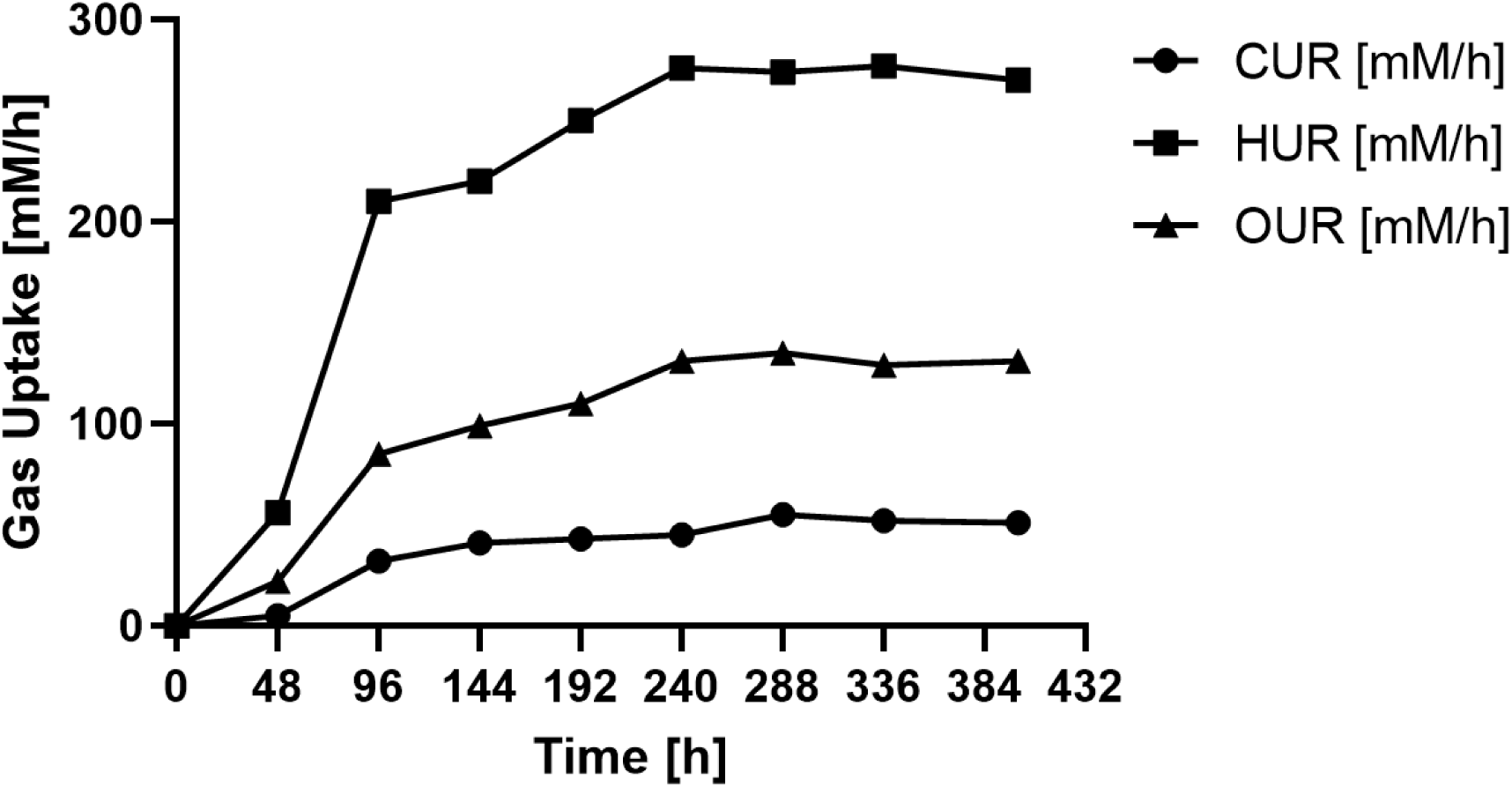
Gas uptake rates for the continuous gas fermentation producing PDC from TPA using the H16 TAPDO *ligABC* strain

### Supplementary Tables

**Table S4:**
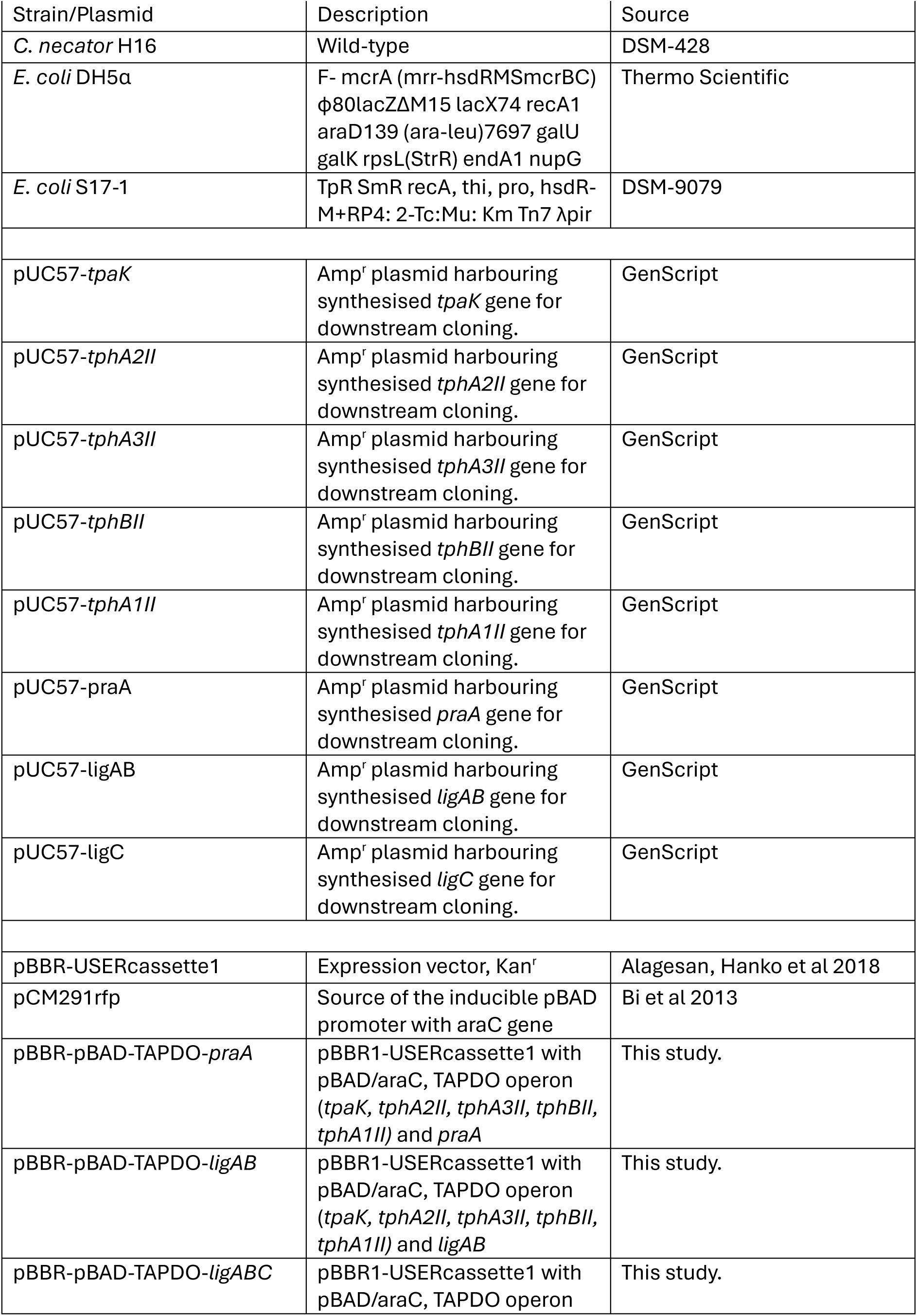
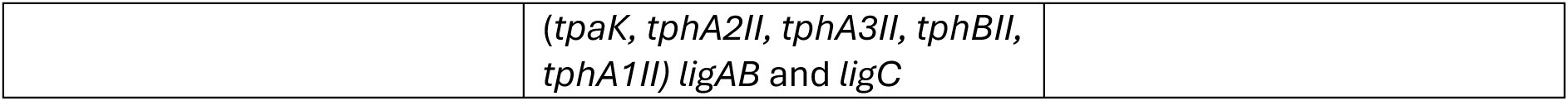
Strains and plasmids used in this study.

**Table S5:**
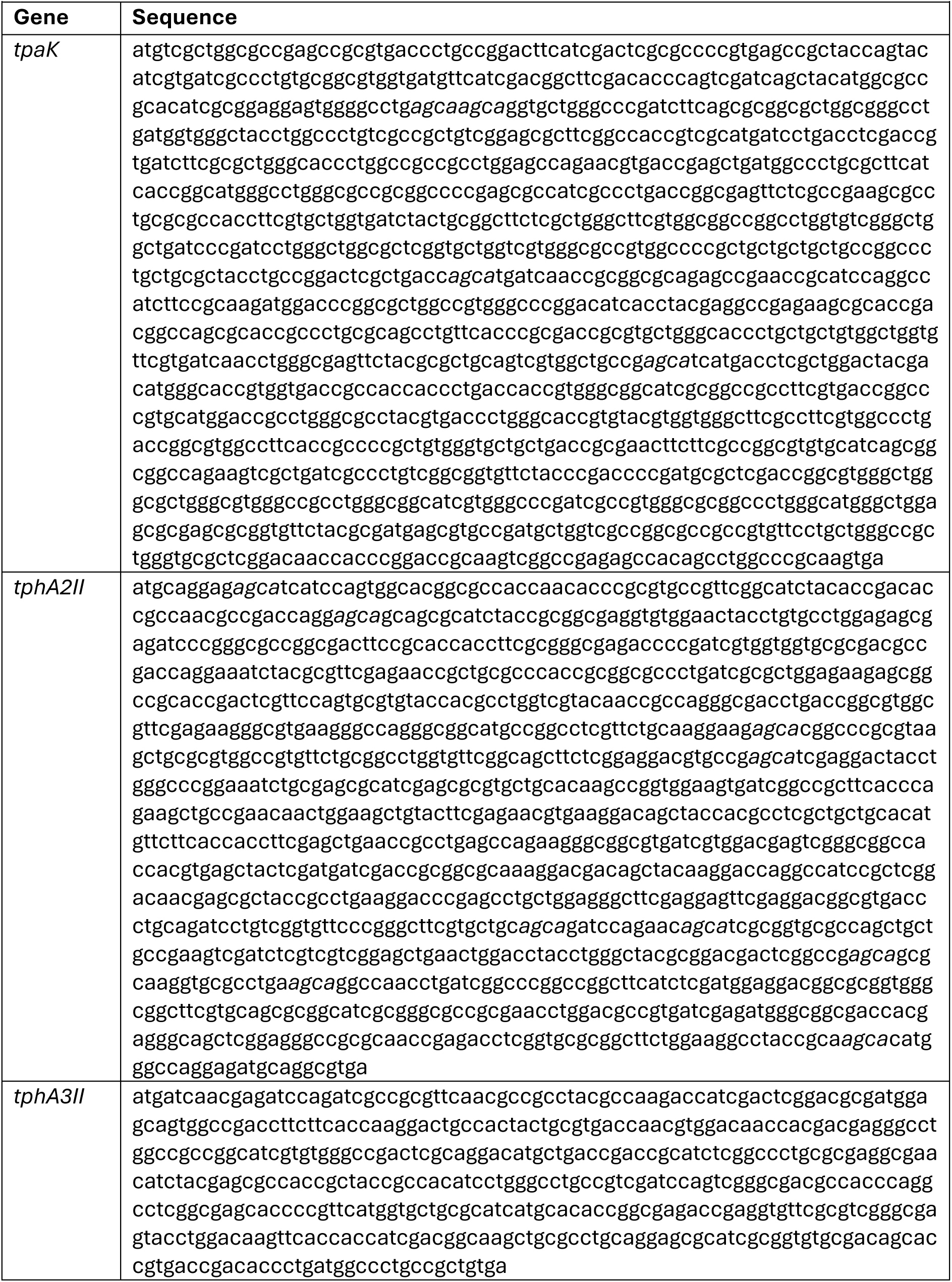
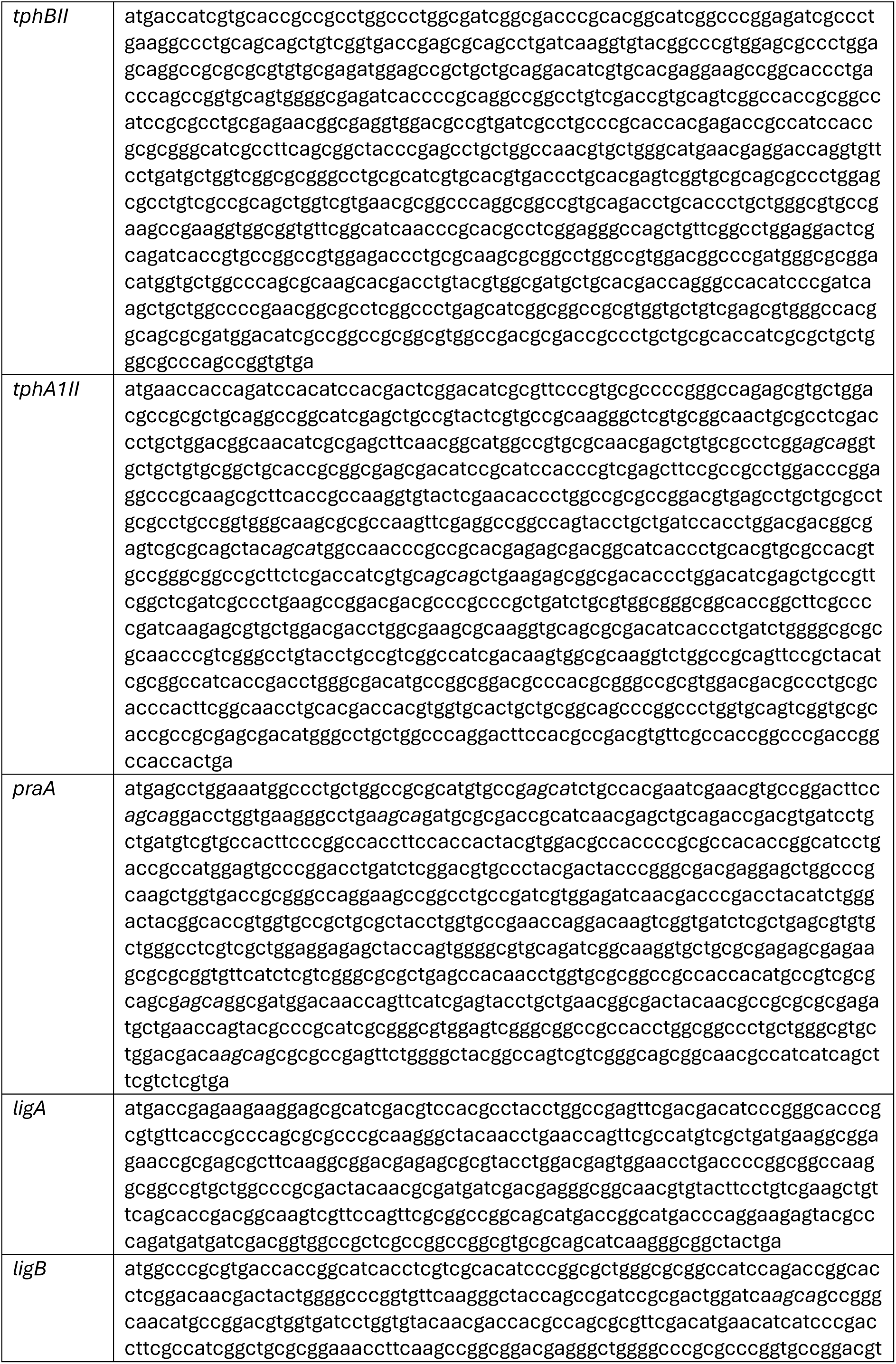
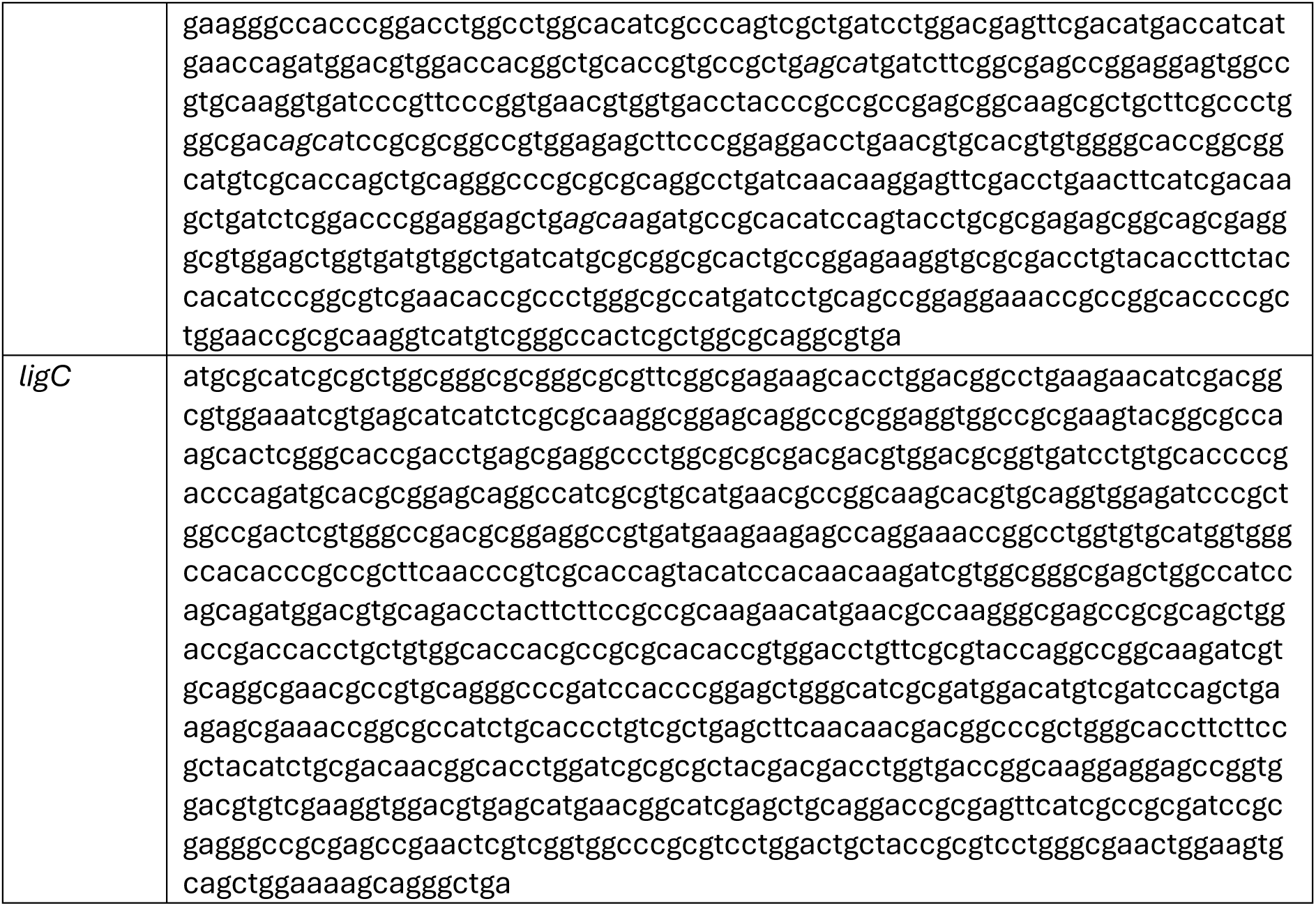
Codon-optimised synthesised genes used in this study.

**Table S6:**
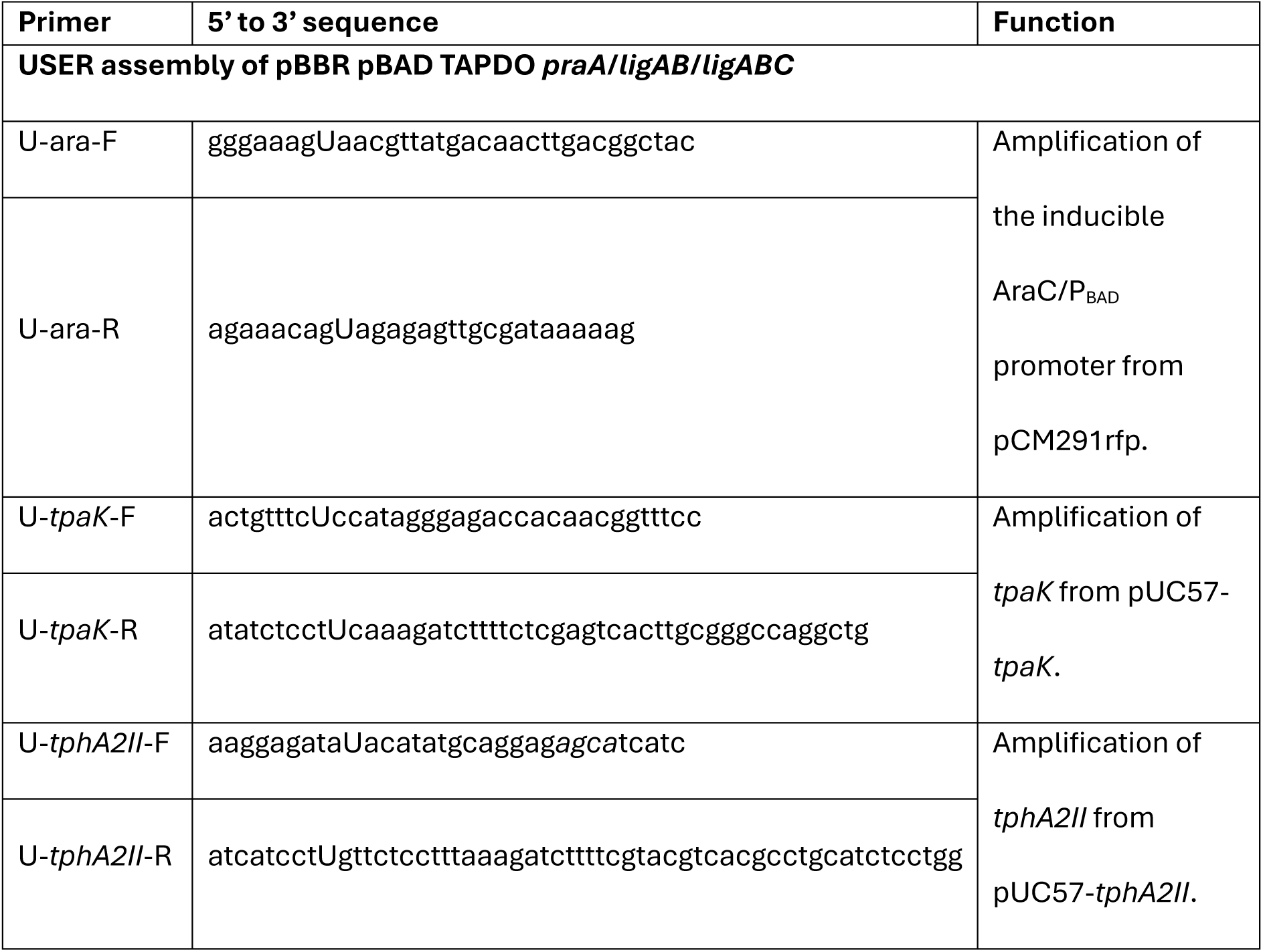
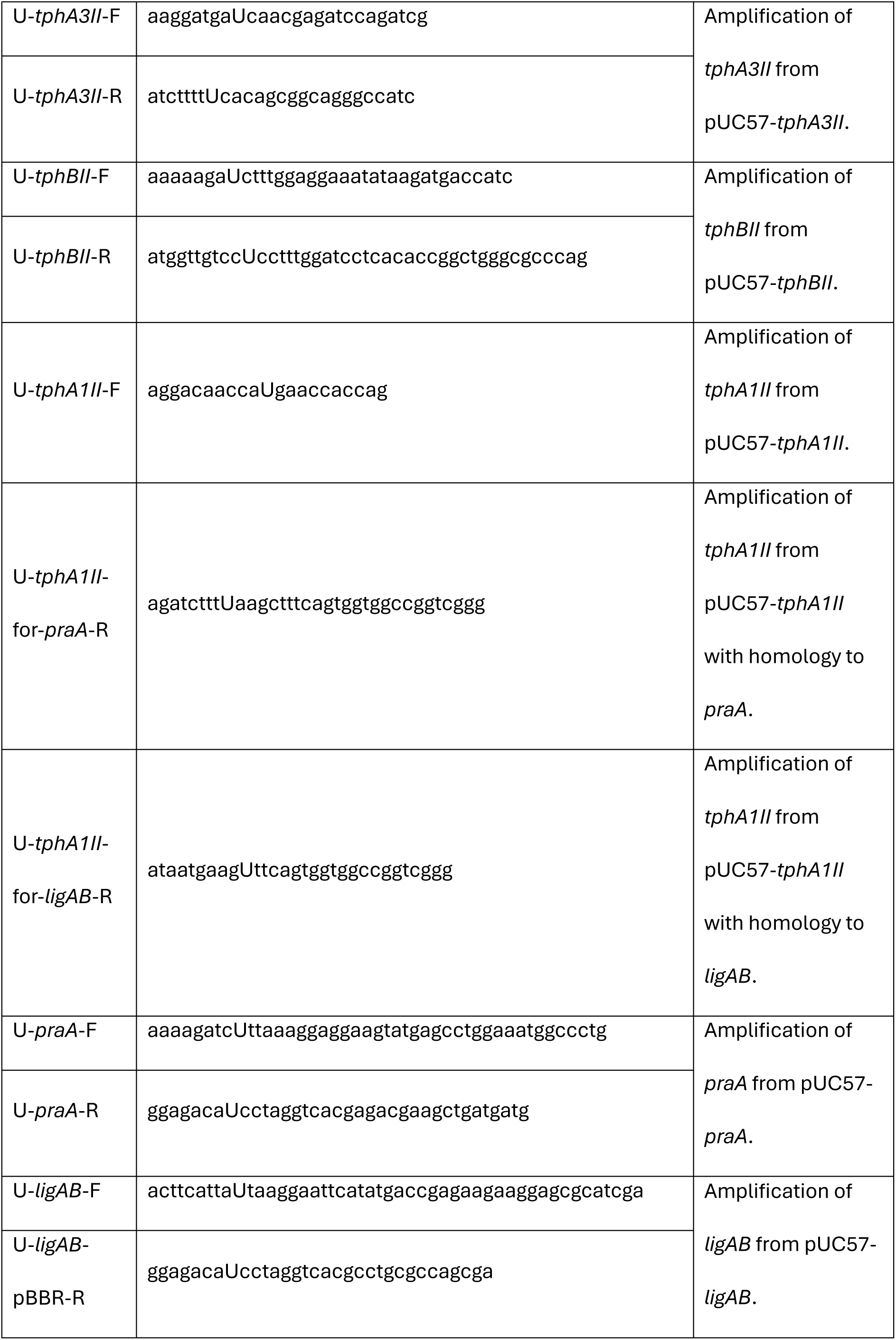
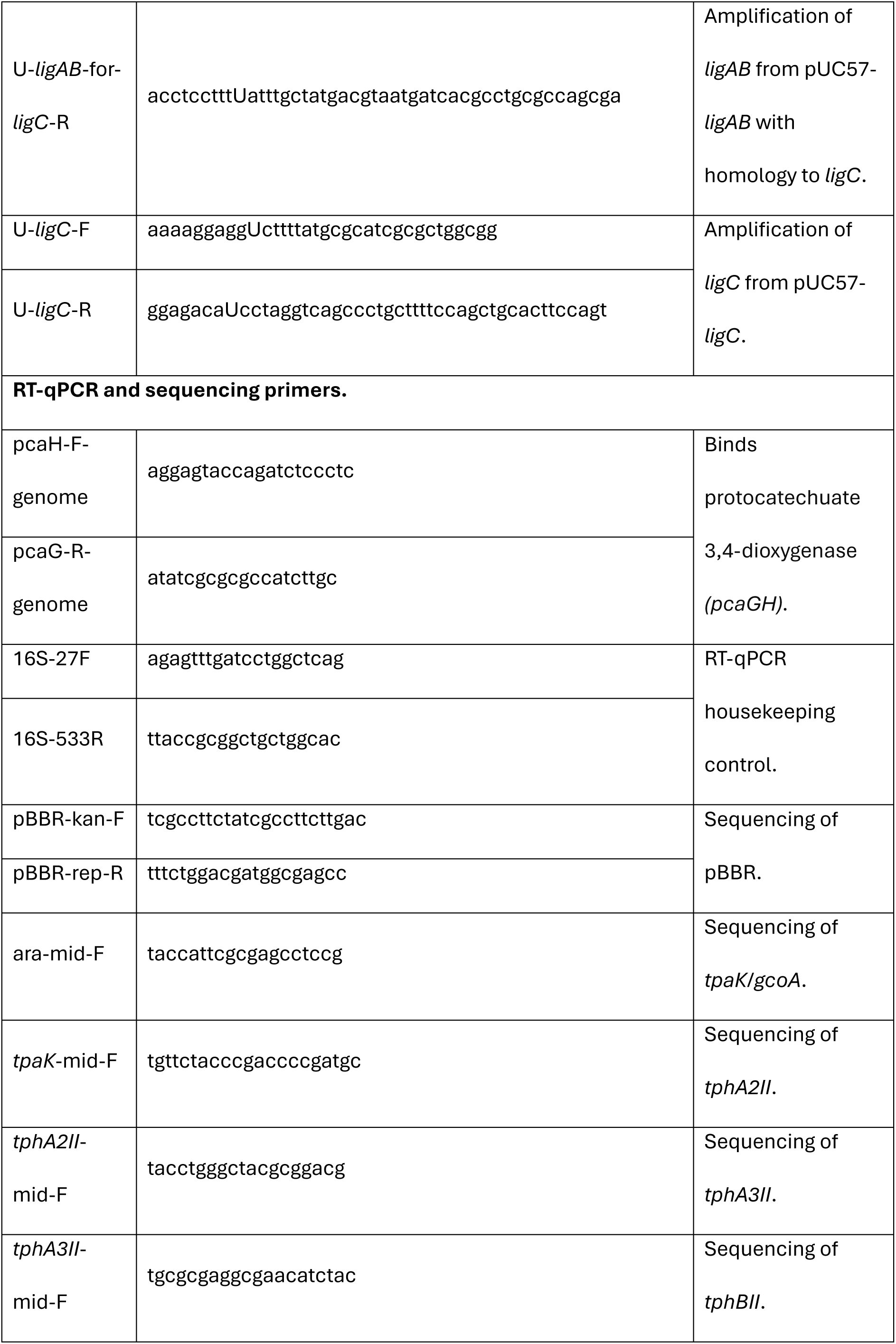
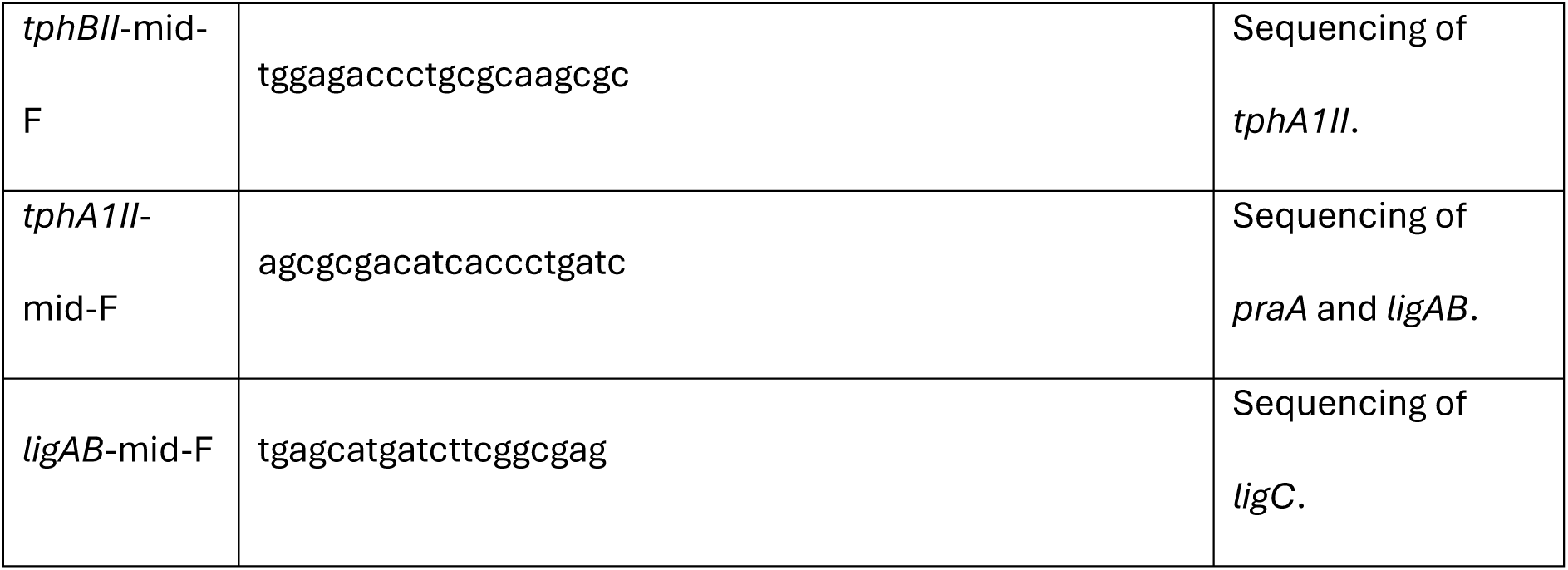
Primers used in this study.

## References

1. Liew FE, Nogle R, Abdalla T, Rasor BJ, Canter C, Jensen RO, et al. Addendum: Carbon-negative production of acetone and isopropanol by gas fermentation at industrial pilot scale. Nature biotechnology. 2025;43(8):1385–7; doi: 10.1038/s41587-025-02767-w.

2. 2. Moore M, Liu VZ, Yang C-K, Cowden Z, Simpson SD. Chapter 18 - Anaerobic gas fermentation: A carbon-refining process for the production of sustainable fuels, chemicals, and food. In: Tuli D, Kasture S, Kuila A, editors. Advanced Biofuel Technologies. Elsevier; 2022. p. 457–74.

3. Santolin L, Riedel SL, Brigham CJ. Synthetic biology toolkit of Ralstonia eutropha (Cupriavidus necator). Applied microbiology and biotechnology. 2024;108(1):450; doi: 10.1007/s00253-024-13284-2.

4. Rodgers S, Conradie A, King R, Poulston S, Hayes M, Bommareddy RR, et al. Reconciling the Sustainable Manufacturing of Commodity Chemicals with Feasible Technoeconomic Outcomes. Johnson Matthey Technology Review. 2021;65(3):375–94; doi: 10.1595/205651321X16137377305390.

5. Bommareddy RR, Wang Y, Pearcy N, Hayes M, Lester E, Minton NP, et al. A Sustainable Chemicals Manufacturing Paradigm Using CO2 and Renewable H2. iScience. 2020;23(6):101218; doi: 10.1016/j.isci.2020.101218.

6. Jawed K, Irorere VU, Bommareddy RR, Minton NP, Kovács K. Establishing Mixotrophic Growth of Cupriavidus necator H16 on CO2 and Volatile Fatty Acids. Fermentation. 2022;8(3):125.

7. Hu P, Chakraborty S, Kumar A, Woolston B, Liu H, Emerson D, et al. Integrated bioprocess for conversion of gaseous substrates to liquids. Proceedings of the National Academy of Sciences of the United States of America. 2016;113(14):3773–8; doi: 10.1073/pnas.1516867113.

8. Ellis LD, Rorrer NA, Sullivan KP, Otto M, McGeehan JE, Román-Leshkov Y, et al. Chemical and biological catalysis for plastics recycling and upcycling. Nature Catalysis. 2021;4(7):539–56; doi: 10.1038/s41929-021-00648-4.

9. Naderi Kalali E, Lotfian S, Entezar Shabestari M, Khayatzadeh S, Zhao C, Yazdani Nezhad H. A critical review of the current progress of plastic waste recycling technology in structural materials. Current Opinion in Green and Sustainable Chemistry. 2023;40:100763; doi: 10.1016/j.cogsc.2023.100763.

10. Satta A, Zampieri G, Loprete G, Campanaro S, Treu L, Bergantino E. Metabolic and enzymatic engineering strategies for polyethylene terephthalate degradation and valorization. Reviews in Environmental Science and Bio/Technology. 2024;23(2):351–83; doi: 10.1007/s11157-024-09688-1.

11. Valenzuela-Ortega M, Suitor JT, White MFM, Hinchcliffe T, Wallace S. Microbial Upcycling of Waste PET to Adipic Acid. ACS Cent Sci. 2023;9(11):2057–63; doi: 10.1021/acscentsci.3c00414.

12. Cho JS, Luo ZW, Moon CW, Prabowo CPS, Lee SY. Metabolic engineering of *Corynebacterium glutamicum* for the production of pyrone and pyridine dicarboxylic acids. Proceedings of the National Academy of Sciences. 2024;121(45):e2415213121; doi: doi:10.1073/pnas.2415213121.

13. Katano A, Mori A, Nonaka D, Mori Y, Noda S, Tanaka T. Biosynthesis of 2,5-pyridinedicarboxylate from glucose via p-aminobenzoic acid in Escherichia coli. Metabolic engineering. 2025;92:252–61; doi: 10.1016/j.ymben.2025.08.011.

14. Gomez-Alvarez H, Iturbe P, Rivero-Buceta V, Mines P, Bugg TDH, Nogales J, et al. Bioconversion of lignin-derived aromatics into the building block pyridine 2,4-dicarboxylic acid by engineering recombinant Pseudomonas putida strains. Bioresource technology. 2022;346:126638; doi: 10.1016/j.biortech.2021.126638.

15. Spence EM, Calvo-Bado L, Mines P, Bugg TDH. Metabolic engineering of Rhodococcus jostii RHA1 for production of pyridine-dicarboxylic acids from lignin. Microbial cell factories. 2021;20(1):15; doi: 10.1186/s12934-020-01504-z.

16. Otsuka Y, Araki T, Suzuki Y, Nakamura M, Kamimura N, Masai E. High-level production of 2-pyrone-4,6-dicarboxylic acid from vanillic acid as a lignin-related aromatic compound by metabolically engineered fermentation to realize industrial valorization processes of lignin. Bioresource technology. 2023;377:128956; doi: 10.1016/j.biortech.2023.128956.

17. Kang MJ, Kim HT, Lee M-W, Kim K-A, Khang TU, Song HM, et al. A chemo-microbial hybrid process for the production of 2-pyrone-4,6-dicarboxylic acid as a promising bioplastic monomer from PET waste. Green Chemistry. 2020;22(11):3461–9; doi: 10.1039/D0GC00007H.

18. Zishuai Wang GX, Yifan Lu and Haijia Su Upcycling of Waste Poly(ethylene terephthalate) into 2,4-Pyridine Dicarboxylic Acid by a Tandem Chemo-Microbial Process. Green Chem Technol. 2025;2(10008); doi: 10.70322/gct.2024.10008.

19. Wu F, Wang S, Zhou D, Gao S, Song G, Liang Y, et al. Metabolic engineering of Escherichia coli for high-level production of the biodegradable polyester monomer 2-pyrone-4,6-dicarboxylic acid. Metabolic Engineering. 2024;83:52–60; doi: 10.1016/j.ymben.2024.03.003.

20. Jones SW, Fast AG, Carlson ED, Wiedel CA, Au J, Antoniewicz MR, et al. CO2 fixation by anaerobic non-photosynthetic mixotrophy for improved carbon conversion. Nature communications. 2016;7(1):12800; doi: 10.1038/ncomms12800.

21. Delmulle T, Bovijn S, Deketelaere S, Castelein M, Erauw T, D’Hooghe M, et al. Engineering Comamonas testosteroni for the production of 2-pyrone-4,6-dicarboxylic acid as a promising building block. Microb Cell Fact. 2023;22(1):188; doi: 10.1186/s12934-023-02202-2.

22. Singh A, Rorrer NA, Nicholson SR, Erickson E, DesVeaux JS, Avelino AFT, et al. Techno-economic, life-cycle, and socioeconomic impact analysis of enzymatic recycling of poly(ethylene terephthalate). Joule. 2021;5(9):2479–503; doi: 10.1016/j.joule.2021.06.015.

23. Schlegel HG, Lafferty R. Growth of ‘Knallgas’ Bacteria (Hydrogenomonas) using Direct Electrolysis of the Culture Medium. Nature. 1965;205(4968):308–9; doi: 10.1038/205308b0.

